# Autophagy-driven Presynaptic Reorganization as a Molecular Signature of Brain Resilience

**DOI:** 10.1101/2025.07.09.663822

**Authors:** David Toppe, Sheng Huang, Janine Lützkendorf, Fan Liu, Marta Maglione, Stephan J. Sigrist

## Abstract

Neural circuits must remain functionally stable while responding flexibly to changing demands, stressors, and aging-related decline. While this balance is thought to be maintained through plasticity programs that integrate molecular, metabolic, and activity-dependent signals to reconfigure synapses structurally and functionally, direct mechanistic models of how such adaptations are orchestrated remain scarce.

Here, we show that targeted impairment of autophagy in the *Drosophila* mushroom body (MB), a key sleep-regulatory and integrative center in the fly brain, triggers a brain-wide remodeling at presynaptic active zones (AZ). Quantitative proteomics revealed a specific upregulation of AZ scaffold proteins (including BRP, RIM, and Unc13A), accompanied by reduced levels of calcium channel subunits and increased Shaker-type potassium channels. These changes occurred largely independent of transcription and highlight a coordinated, excitability-tuning response centered on the AZ. Behaviorally, MB-specific autophagy impairment increased sleep and modestly extended lifespan. These adaptations resembled a previously described resilience program termed PreScale, which promotes restorative sleep homeostasis in response to sleep deprivation and early, still reversible brain aging. Conversely, overexpression of *Atg5* in the MB delayed the onset of PreScale. Notably, autophagic disruption confined to MB neurons also caused widespread, non-cell autonomous accumulation of Ref(2)P and ATG8a-positive aggregates across the brain, revealing systemic propagation of proteostatic stress.

Together, our findings identify MB autophagy as a key regulator of synaptic architecture and sleep-associated resilience. Such early acting programs may actively preserve circuit function and behavioral output by regulating synaptic plasticity, and define a genetically tractable model for how local stress signals can orchestrate brain-wide adaptation via post-transcriptional synaptic reprogramming.

## Introduction

Neuronal networks are continuously challenged by fluctuating metabolic conditions ^1,2^, insufficient sleep ^3–5^, and the gradual loss of proteostasis during aging ^6,7^. While these stressors threaten cellular and circuit stability, the brain has evolved mechanisms to maintain function in the face of adversity. Indeed, many organisms display a remarkable capacity for brain resilience well into advanced mid-life, suggesting that adaptive responses are engaged prior to the onset of overt degeneration ^6,8–11^. Such early acting programs may actively preserve circuit function and behavioral output through regulated synaptic plasticity and organization.

Autophagy, a lysosome-mediated degradation pathway that clears damaged or superfluous cytoplasmic components, is central to neuronal health and lifespan ^12–16^. Autophagic efficiency declines with age ^17–21^ and is impaired in numerous neurodegenerative ^22–25^ and neurodevelopmental conditions ^26,27^. In neurons, autophagy is not only essential for maintaining local proteostasis, particularly at long-lived presynaptic terminals ^15,28,29^, but may also participate in broader plasticity processes ^30^. Yet, how local autophagy dysfunction might influence global circuit remodeling and resilience remains poorly understood.

Sleep has emerged as a powerful modulator of neuronal plasticity and resilience. Sleep deprivation perturbs cellular homeostasis and impairs cognitive performance ^3,31^, whereas restorative sleep promotes synaptic tuning ^32–34^ and metabolic waste clearance from the brain ^35^. In *Drosophila*, the mushroom body (MB), a higher brain center for integration of multisensory information and metabolic signals ^36^, has been identified as a key regulator of sleep homeostasis ^37,38^, integrating both sleep-promoting and arousal-related inputs ^39^. The MB is thus well positioned to coordinate brain-wide responses to internal stress.

We previously showed that local autophagy impairment within the MB intrinsic Kenyon cells (KCs) is sufficient to trigger a brain-wide increase of the active zone (AZ) protein Bruchpilot (BRP) ^40^, a phenotype which also represents a key feature of the brain-wide presynaptic remodeling program termed PreScale ^11^. PreScale shows features of a resilience-promoting plasticity state that is typically induced in response to sleep deprivation ^41–43^ or emerges progressively during early aging ^11,44^ and its activation has been linked to improved stress tolerance and survival ^11^. Indeed, protective interventions such as dietary spermidine ^44^ or enhanced sleep ^11^ delay or reduce PreScale activation, supporting the idea that it represents a compensatory response to accumulating physiological stress. In the present study, we build on these observations and provide a detailed molecular and functional characterization of PreScale, identifying its structural logic, temporal dynamics, and mechanistic links to resilience under conditions of local autophagy impairment.

Using RNAi-based approaches to impair autophagy function specifically within MB KCs, we show that local disruption of proteostasis inside the MB is sufficient to accelerate the activation of PreScale across the brain during early aging. Despite local autophagy failure, affected animals exhibit preserved or even enhanced sleep and survival, indicating that this AZ remodeling response is not merely compensatory but functionally protective. Quantitative proteomic analysis reveals that this response is characterized by a coordinated increase in AZ scaffold proteins, including BRP, RIM, Liprin-α, and Unc13A, alongside reduced abundance of voltage-gated calcium channels (VGCCs) and a concomitant upregulation of the voltage-gated potassium channels Shaker and Shaw. These combined changes likely shift presynaptic excitability and synaptic vesicle release probability in a manner that supports robust circuit function under stress.

Together, our findings suggest that PreScale constitutes a modular, resilience-promoting synaptic plasticity program, that can be locally initiated by autophagy dysfunction in the MB, but is deployed across the brain to recalibrate synaptic plasticity and AZ organization in order to preserve function in face of imbalanced proteostasis.

## Results

A transient, brain-wide form of presynaptic plasticity — marked by increased abundance of BRP and associated active zone (AZ) components — was previously described and termed PreScale ^11^. PreScale is induced in response to sleep deprivation ^41–43,45^, and its genetic mimicry through increased *brp* gene copy number has been shown to be sufficient to enhance daily sleep ^41^. In parallel, the early phase of *Drosophila* aging (up to ∼20–30 days) is also characterized by a PreScale-like AZ remodeling process, and again, genetic mimicry through increased *brp* gene copy number suggested that this state contributes to organismal resilience and survival ^11,44^. Supporting this view, interventions that promote anti-aging resilience, such as dietary spermidine supplementation or pharmacologically enhanced sleep, were found to delay or reduce the early need to engage PreScale during brain aging ^11,44^. Notably, the difference in PreScale expression and sleep modulation between protected and control animals peaked at around 15 to 20 days of age, highlighting this early time window as a critical phase for resilience programming ^11^.

### MB autophagy bi-directionally regulates brain-wide PreScale during early aging

In terms of brain regions that may be critical for coordinating brain-wide PreScale responses, the *Drosophila* mushroom body (MB) has emerged as a central regulator of sleep homeostasis ^37,38^, integrating both sleep-promoting and wake-promoting circuits and signals ^39^. But what kinds of molecular signals does the MB process to fulfill this integrative role? In this regard, we previously found that the impairment of autophagy specifically within the MB-intrinsic Kenyon cells (KCs) triggered a brain-wide increase in BRP levels, a hallmark of PreScale, when examined during early aging at 10d of age ^40^.

Given the progressive and dynamically modulable build-up of PreScale during early aging in wild-type flies ^11,44^, we started by systematically investigating the kinetics of PreScale activation in response to MB-specific autophagy manipulation across the early aging period between day 5 and day 30 of age (Fig. 1). In animals with early MB-specific *Atg5* knockdown under control of the strong and KC-specific *vt30559*-Gal4 driver ^46^, BRP levels were significantly elevated compared to wildtype *vt30559*-Gal4/+ controls both at 5 (Fig. 1A,A’) and 15 days (Fig. 1B,B’) after eclosion. Notably, by 30 days of age — marking the end of the early aging period — BRP levels in MB-autophagy-impaired animals and control flies were indistinguishable (Fig. 1C,C’). This early-onset BRP upregulation suggests that PreScale, which typically builds up gradually during aging ^11^, exhibits accelerated activation when autophagy is impaired within the MB, before reaching a normal age-matched end-state by 30 days. Notably, in the converse experiment, the genetic overexpression of *Atg5* specifically within the MB KCs (*vt30559*-Gal4>UAS-*Atg5*-GFP) led to a specific, significant reduction in brain-wide BRP levels once again at 15 days (Fig. 1E,E’), but not anymore at 30 days (Fig. 1F,F’) or at 5 days of age (Fig. 1D,D’). These findings support a model in which the functional status of MB autophagy bi-directionally modulates the timing of activation of brain-wide PreScale-related AZ-remodeling during early *Drosophila* aging (Fig. 1G), highlighting its instructive role in resilience-linked synaptic plasticity.

**Fig. 1.**
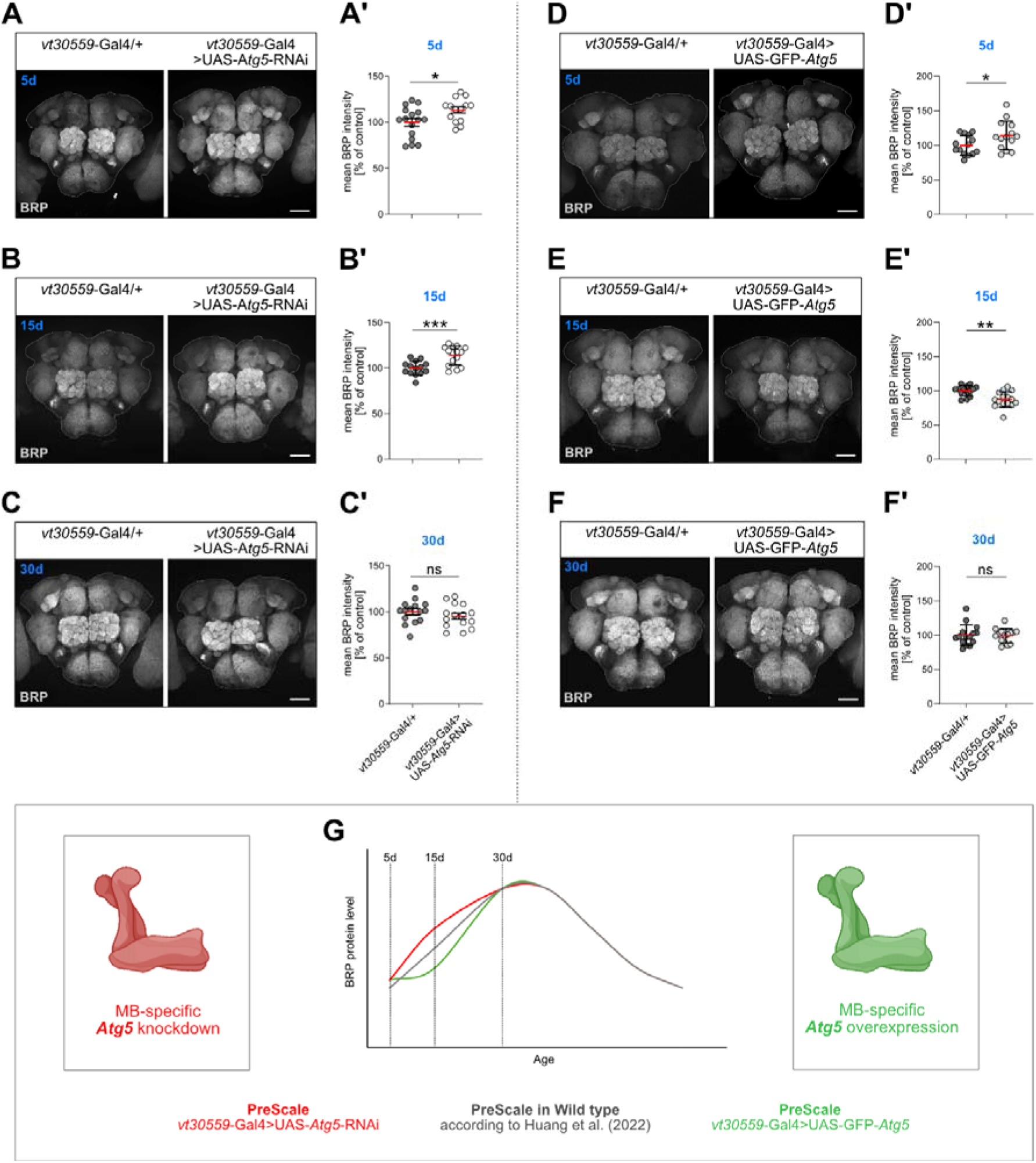
Autophagy within the MB controls brain-wide presynaptic plasticity (PreScale) during early aging. **(A-C)** Representative confocal max projections of adult brains of *vt30559*-Gal4>UAS-*Atg5*-RNAi and control *vt30559*-Gal4/+ female flies at 5d, 15d or 30d of age. Brains were immunostained against BRP^Nc82^, and measured whole brain mean BRP intensities for all groups of a specific age were normalized to the respective *vt30559*-Gal4/+ control group. **(A)** Representative confocal max projections and **(A’)** quantification of BRP intensities for 5d old female flies. Scale bar = 50 μm; n = 14-16; Two-tailed t-test **(B)** Representative confocal max projections and **(B’)** quantification of BRP intensities for 15d old female flies. Scale bar = 50 μm; n = 7-16; Two-tailed t-test **(C)** Representative confocal max projections and **(C’)** quantification of BRP intensities for 30d old female flies. Scale bar = 50 μm; n = 11-15; Two-tailed t-test. **(D-F)** Representative confocal max projections of adult brains of *vt30559*-Gal4>UAS-GFP-*Atg5* and control *vt30559*-Gal4/+ female flies at 5d, 15d or 30d of age. Brains were immunostained against BRP^Nc82^, and measured whole brain mean BRP intensities for all groups of a specific age were normalized to the respective *vt30559*-Gal4/+ control group. **(D)** representative confocal max projections and **(D’)** quantification of BRP intensities for 5d old female flies. Scale bar = 50 µm; n = 14; two-tailed t-test. **(E)** representative confocal max projections and **(E’)** quantification of BRP intensities for 15d old female flies. Scale bar = 50 µm; n = 14-15; two-tailed t-test. **(F)** representative confocal max projections and **(F’)** quantification of BRP intensities for 30d old female flies. Scale bar = 50 µm; n = 15; two-tailed t-test. **(G)** Model of age-dependent increase of brain-wide BRP levels as a surrogate of PreScale activation in wildtype flies and upon genetic manipulation of MB autophagy. * = p<0.05; ** = p<0.01; *** = p<0.001; **** = p<0.0001; ns, not significant. Data presented as mean ± SD.

### MB autophagy bi-directionally controls resilience-associated sleep patterns during early aging

Considering the MB’s central role in sleep regulation ^37–39^ and our observation that early MB-specific autophagy impairment triggers accelerated brain-wide PreScale-like synaptic remodeling (Fig. 1A-C), we asked whether homeostatic sleep control during early aging might be sensitive to proteostatic disruptions within the MB. Indeed, 15-day-old female flies with MB-specific *Atg5* knockdown (*vt30559*-Gal4>UAS-*Atg5*-RNAi) exhibited significantly increased total amounts of daily sleep, including both daytime and nighttime sleep, compared to age- and sex-matched controls (Fig. 2A,B). Moreover, *Atg5* knockdown in the MB also enhanced nighttime sleep in the context of the chronic short-sleeping mutant sleepless (*sss^P1^*; *vt30559*-Gal4>UAS-*Atg5*-RNAi), which served as a sensitized genetic background (Fig. 2C,D). This result places MB autophagy function upstream of a known core sleep regulator and suggests that proteostasis within the MB contributes to sleep control at a high hierarchical level of the sleep homeostat.

**Fig. 2.**
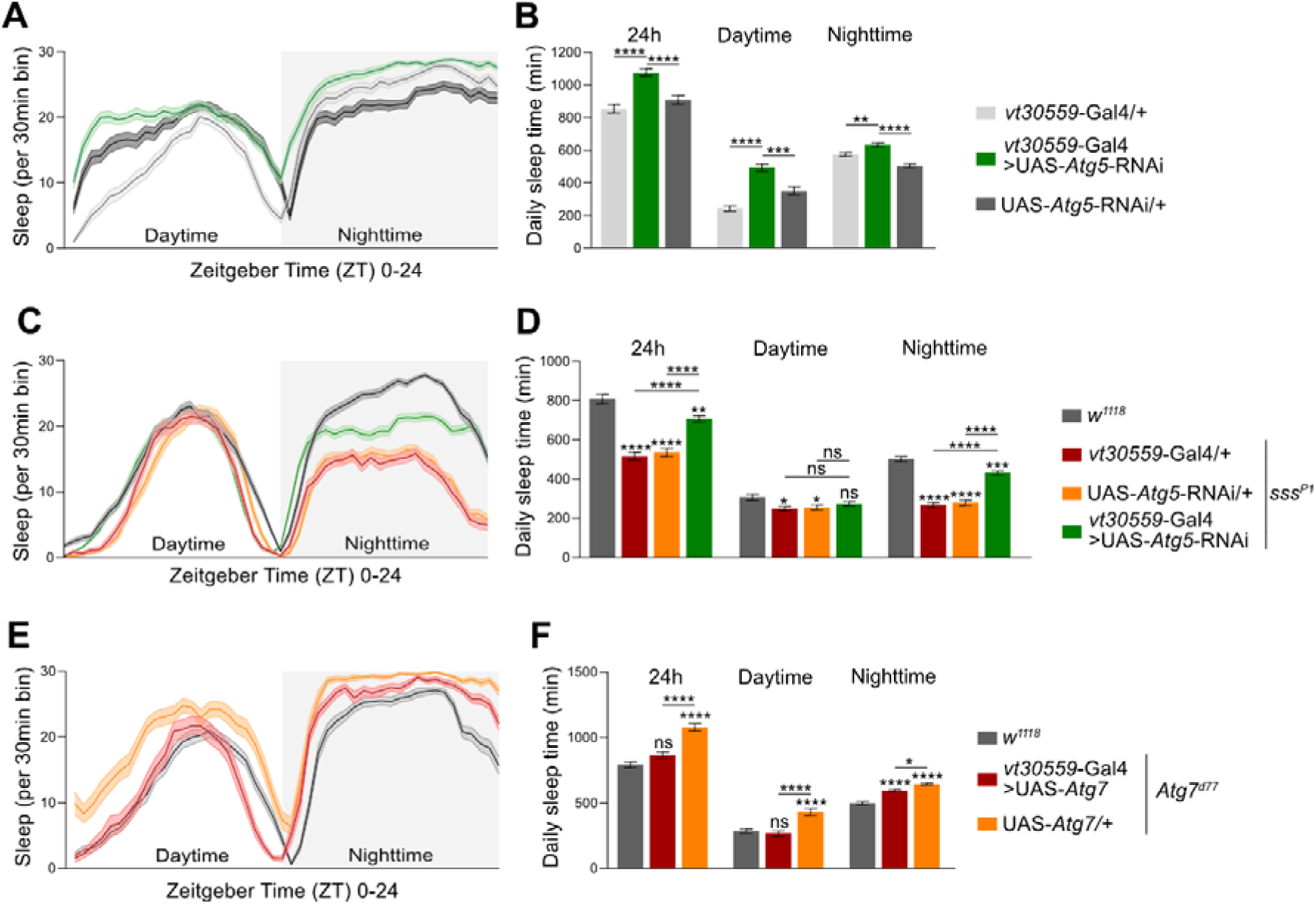
The functional status of MB-intrinsic autophagy bi-directionally controls sleep homeostasis. **(A and B)** Sleep parameters of 15d old female flies with MB-specific autophagy impairment (*vt30559*-Gal4>UAS-*Atg5*-RNAi) compared to two age- and sex-matched controls carrying only one of the two components of the binary Gal4/UAS system (*vt30559*-Gal4/+ and UAS-*Atg5*-RNAi/+). Sleep data was acquired and averaged over 3 consecutive days. **(A)** Average sleep profiles over 24h (Zeitgeber Time 0-24) plotted in 30 mins bins, including 12h light (ZT0-ZT12, Daytime) and 12h darkness (ZT12-ZT24, Nighttime). **(B)** Averaged daily sleep amounts in mins over 24h, during Daytime and Nighttime. n = 45-46. **(C and D)** Sleep parameters of 5d old female *sss^P1^* mutants with MB-restricted knockdown of *Atg5* gene expression (*sss^P1^;vt30559*-Gal4>UAS-*Atg5*-RNAi) were compared to age- and sex-matched controls in the same *sss^P1^* genetic background (*sss^P1^*;UAS-*Atg5*-RNAi/+ and *sss^P1^;vt30559*-Gal4/+) as well as to a *w^1118^* wild-type control. Sleep data was acquired and averaged over 3 consecutive days. **(C)** Average sleep profiles over 24h (Zeitgeber Time 0-24) plotted in 30 mins bins, including 12h light (ZT0-ZT12, Daytime) and 12h darkness (ZT12-ZT24, Nighttime). **(D)** Averaged daily sleep amounts in mins over 24h, during Daytime and Nighttime. n = 46-58. **(E and F)** Sleep parameters of 5d old female *Atg7^d77^* mutant flies with MB-specific expression of *Atg7* (*Atg7^d77^;vt30559*-Gal4>UAS-*Atg7*), as well as of *Atg7^d77^*,UAS-*Atg7*/+ genetic controls and *w^1118^*-type flies. Sleep data was acquired and averaged over 3 consecutive days. **(E)** Average sleep profiles over 24h (Zeitgeber Time 0-24) plotted in 30 mins bins, including 12h light (ZT0-ZT12, Daytime) and 12h darkness (ZT12-ZT24, Nighttime). **(F)** Averaged daily sleep amounts in mins over 24h, during Daytime and Nighttime. n = 22-45. * = p<0.05; ** = p<0.01; *** = p<0.001; **** = p<0.0001; ns, not significant, one-way ANOVA with Tukey’s post-hoc test for multiple comparisons. Data presented as mean ± SEM.

To test whether restoring autophagy selectively within the MB is sufficient to influence sleep behaviour, we examined isogenized *Atg7* null mutants (*Atg7^d77^*), which systemically lack autophagy ^47^. Consistent with prior reports showing increased sleep upon global loss of neuronal autophagy ^48^, *Atg7^d77^* null mutants showed robust sleep elevation relative to wildtype *w^1118^* (Fig. 2E,F). Remarkably, re-expression of *Atg7* specifically within the MB KCs (*Atg7^d77^*;*vt30559*-Gal4>UAS-*Atg7*) significantly reduced both daytime and nighttime sleep toward wild-type sleep levels (Fig. 2E,F).

Together, these findings demonstrate that MB autophagy exerts bidirectional control over sleep regulation during early aging and highlight its instructive role in a resilience-associated behavioral program.

### MB-specific knockdown of *Atg5* modestly enhances longevity and preserves oxidative stress resistance

We wondered whether the MB-triggered resilience response—specifically, the accelerated onset of PreScale during early aging (Fig. 1A-C) and the associated changes in sleep patterns upon MB-specific autophagy impairment (Fig. 2A,B)—might be able to compensate for the inherently detrimental loss of autophagic degradation and proteostasis inside the MB. Loss of autophagy is *per se* expected to shorten lifespan and to increase the sensitivity to additional oxidative stress ^16,47,49–51^. As anticipated, pan-neuronal knockdown of *Atg5* (*elav*-Gal4>UAS-*Atg5*-RNAi) indeed significantly reduced lifespan in both male and female flies compared to genetic controls (Fig. 3A,B).

**Fig. 3.**
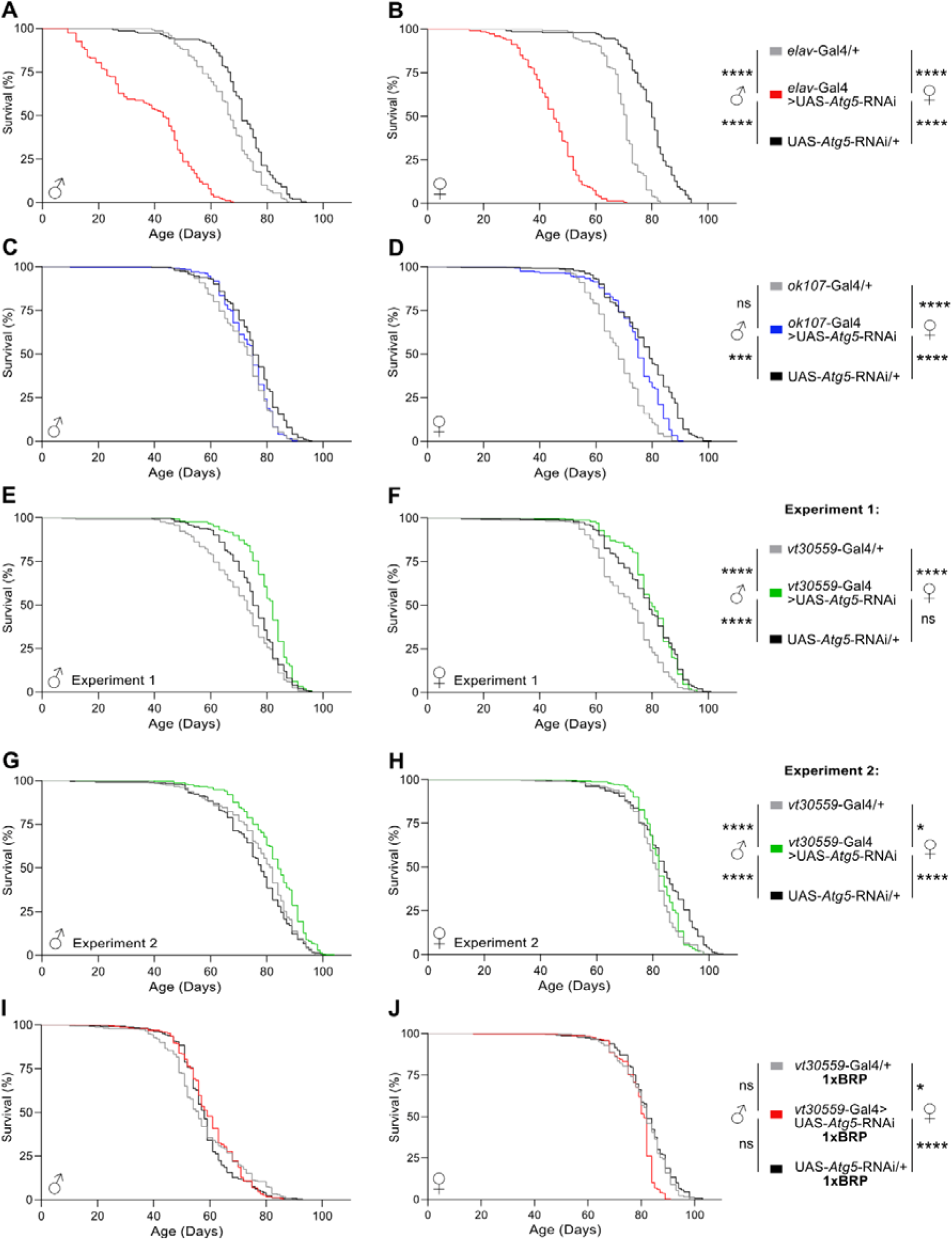
Improved survival of flies with MB-specific *Atg5* knockdown requires adequate BRP levels. **(A and B)** Kaplan-Meier survival curves of male **(A)** and **(B)** female flies with pan-neuronal *Atg5* knockdown (*elav*-Gal4>UAS-*Atg5*-RNAi, red curve) compared to *elav*-Gal4/+ and UAS-*Atg5*-RNAi/+ control flies. n = 144-148 per genotype and sex. **(C and D)** Kaplan-Meier survival curves of male **(C)** and **(D)** female flies with MB-specific *Atg5* knockdown (*ok107*-Gal4>UAS-*Atg5*-RNAi, blue curve) compared to *ok107*-Gal4/+ and UAS-*Atg5*-RNAi/+ control flies. n = 242-247 per genotype and sex. **(E and F)** Kaplan-Meier survival curves of male **(E)** and **(F)** female flies with MB-specific *Atg5* knockdown (*vt30559*-Gal4>UAS-*Atg5*-RNAi, green curve) compared to *vt30559*-Gal4/+ and UAS-*Atg5*-RNAi/+ control flies. n = 240-248 per genotype and sex. The UAS-*Atg5*-RNAi/+ controls in (C) and (D) are the same as in (E) and (F). **(G and H)** Kaplan-Meier survival curves of male **(G)** and female **(H)** *vt30559*-Gal4>UAS-*Atg5*-RNAi flies compared to *vt30559*-Gal4/+ and UAS-*Atg5*-RNAi/+ controls. Experiment performed with independent cohorts of flies from the ones used in E + F. n = 236-254 per genotype and sex. Pairwise Log-rank (Mantel-Cox) test comparison. **(I and J)** Kaplan-Meier survival curves of male **(I)** and female **(J)** flies expressing the same combinations of transgenes as the genotypes in G and H, but additionally carrying the *Brp^c04298^* null allele in heterozygosity to reduce *Brp* gene dosage by 50% (referred to as 1xBRP). n = 234-251 per genotype and sex. Pairwise Log-rank (Mantel-Cox) test comparison. * = p<0.05; ** = p<0.01; *** = p<0.001; **** = p<0.0001; ns, not significant; pairwise Log-rank (Mantel-Cox) test comparison.

In contrast, MB-specific *Atg5* knockdown using the strongly MB-targeting *ok107*-Gal4 driver line ^52^, did not compromise longevity (Fig.3C,D). In fact, we even observed a mild trend toward increased survival in males (Fig. 3C) and a modest but significant extension in females relative to *ok107*-Gal4/+ controls (Fig. 3D), although both sexes remained slightly shorter lived than the UAS-*Atg5*-RNAi/+ controls. Intriguingly, when autophagy was impaired using the strong and highly KC-specific *vt30559*-Gal4 driver line ^53^, median survival of male flies with MB-specific autophagy impairment was even significantly extended by several days compared to both genetic controls (Fig. 3E). Female *vt30559*-Gal4>UAS-*Atg5*-RNAi flies also outlived *vt30559*-Gal4/+ controls and showed a positive survival trend relative to UAS-*Atg5*-RNAi/+ control flies, particularly between day 60 and day 80 of age (Fig. 3F). These findings suggest that local impairment of MB autophagy may engage protective or compensatory responses that support overall organismal survival.

To assess whether MB-specific autophagy impairment also affects systemic stress resilience, we subjected *ok107*-Gal4>UAS-*Atg5*-RNAi and *vt30559*-Gal4>UAS-*Atg5*-RNAi flies, along with age-matched controls, to chronic oxidative stress (2% H□O□ treatment). Despite impaired proteostasis in the central MB, both male and female flies maintained oxidative stress resistance comparable to controls (Extended Data Fig. 1).

Taken together, these findings unexpectedly indicate that MB-specific autophagy impairment results in even slightly enhanced organismal longevity and normal oxidative stress resilience, despite the imbalanced proteostasis in a major brain region. This suggests that local autophagy dysfunction within the central integrative MB structure might be effectively counteracted through adaptive mechanisms, potentially in a brain-wide manner. The transient increase in BRP abundance that we observed during early aging (Fig. 1) raised the possibility that a PreScale-like presynaptic remodeling process may be engaged to mediate these beneficial effects. Building on these observations, we next tested whether interfering with PreScale by reducing *brp* gene dosage would compromise the organismal resilience phenotype of *vt30559*-Gal4>UAS-*Atg5*-RNAi flies (Fig. 3E,F). In independent cohorts of *vt30559*-Gal4>UAS-*Atg5-*RNAi flies and genetic controls, we therefore introduced a heterozygous *brp* null mutation (*Brp^c04298^*) to reduce *brp* gene dosage (1xBRP). Consistent with the first experiment (Fig. 3E,F), male *vt30559*-Gal4>UAS-*Atg5*-RNAi flies with normal wildtype BRP levels (2xBRP) once again showed a significant lifespan extension compared to both controls (Fig. 3G), while females displayed a more modest effect (Fig. 3H). However, this lifespan benefit was completely abolished in males (Fig. 3I), and lifespan was even significantly attenuated in females with MB-specific autophagy impairment in the 1xBRP background (Fig. 3J), indicating that adequate BRP levels are indeed functionally required, at least in part, for promoting survival in light of disrupted MB proteostasis.

Jointly, these findings suggest that MB-autophagy acts as a central regulator of the timing, magnitude, and persistence of brain-wide PreScale responses. Under conditions of autophagy impairment, the MB triggers accelerated and robust PreScale activation during early aging, which promotes resilience and contributes to increased survival. In contrast, preserved MB-autophagy delays PreScale onset, while insufficient BRP dosage prevents sustained PreScale activation and negates its protective effect on survival. These results establish a mechanistic link between local autophagy function within the central MB and organismal resilience, acting through a temporally tuned brain-wide remodeling of presynaptic AZs.

### Coordinated brain-wide remodeling of presynaptic architecture and protein homeostasis in response to MB autophagy impairment

Our results suggest that PreScale functions as a generic, resilience-mediating plasticity program mobilized by the brain in response to diverse physiological challenges. At the molecular level, this state was previously characterized by increased abundance of key active zone (AZ) proteins such as BRP and Unc13A while core components of the vesicle fusion machinery, such as Syntaxin-1, remained largely unchanged ^41^. However, a comprehensive molecular analysis of PreScale has been lacking, and the broader regulatory logic, particularly regarding non-AZ components and the levels at which these changes occur (transcriptional, translational, or post-translational), remained unclear.

To address this, we examined how local, MB-specific autophagy impairment alters the presynaptic proteome on a brain-wide level. We performed label-free quantitative proteomics on hand-dissected adult brains from *ok107*-Gal4>UAS-*Atg5*-RNAi flies and compared them to *ok107*-Gal4/+ controls. Across four biological replicates per genotype, we detected approximately 3,500 individual proteins (present in ≥3 of 4 replicates). Of these, 471 proteins were significantly upregulated, and 172 significantly downregulated in response to MB-specific *Atg5* knockdown.

Subsequent unbiased Gene Ontology (GO) functional enrichment analysis using Metascape ^54^ revealed a strong enrichment of endoplasmic reticulum (ER)-associated categories among upregulated protein hits (Fig. 4A,C), including the GO terms “endoplasmic reticulum” (GO:0005783), “rough ER” (GO:0005791), “response to ER stress” (GO:0034976), and “protein folding” (GO:0006457). Protein–protein interaction (PPI) analysis further confirmed that ER and endomembrane-associated proteins formed distinct and coherent functional modules (“endomembrane system“; Fig. 4E).

**Fig. 4.**
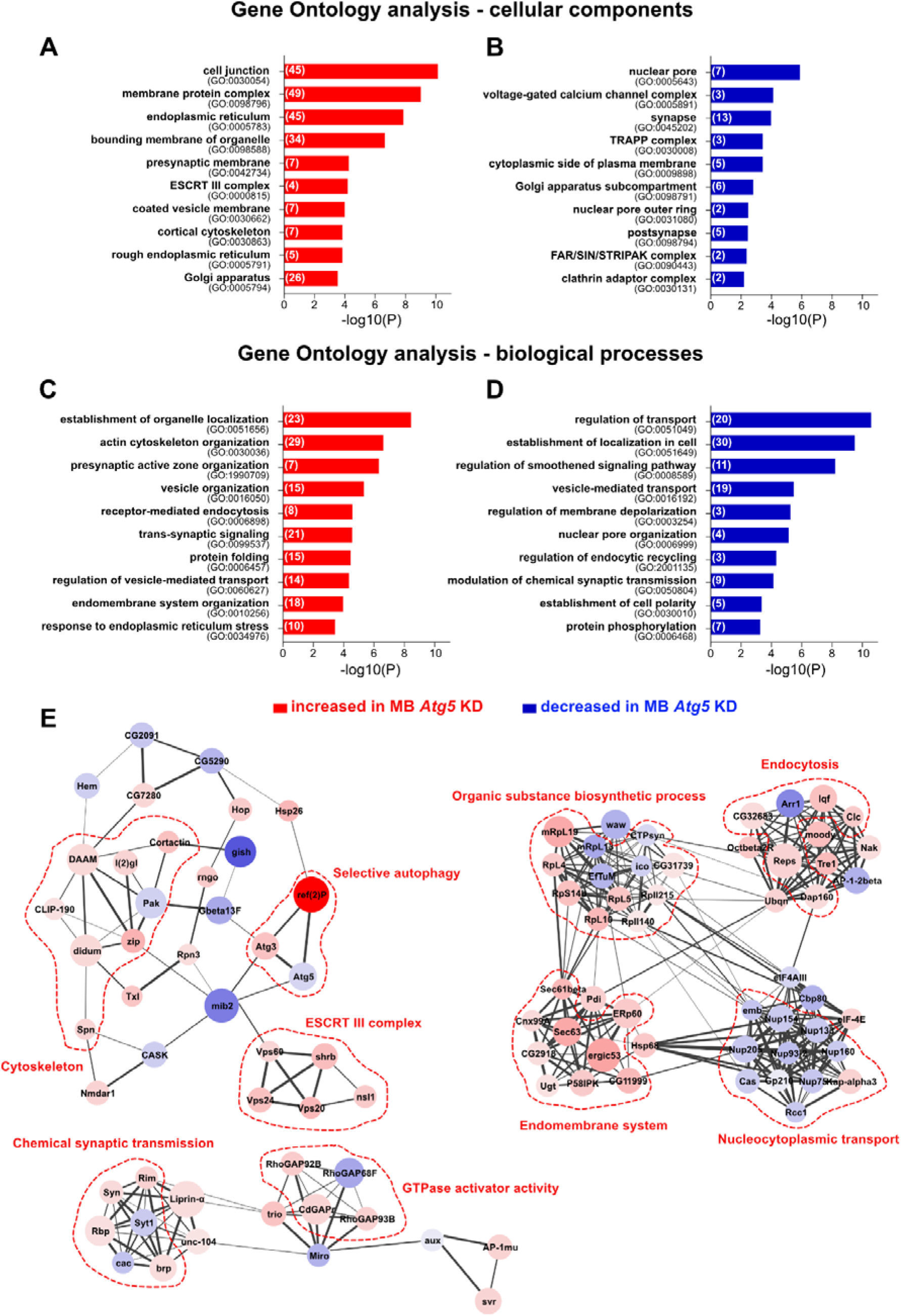
Functional enrichment analysis and interactome of significantly changed proteins in fly brains with MB-specific *Atg5* knockdown. Functional GO enrichment analysis was performed with Metascape ^54^. **(A and B)** Selected enriched ‘cellular components’ GO terms overrepresented in fly brains with MB-specific knockdown of *Atg5* (*ok107*-Gal4>UAS-*Atg5*-RNAi) (**A**, increased) or in the genetic control (*ok107*-Gal4/+) (**B**, decreased). The number of significantly changed proteins identified for each GO term is indicated in brackets. A single protein can be classified into multiple different GO terms. **(C and D)** Selected enriched ‘biological processes’ GO terms overrepresented in flies with MB-specific knockdown of *Atg5* (*ok107*-Gal4>UAS-*Atg5*-RNAi) (**C**, increased) or in the wild-type control (*ok107*-Gal4/+) (**D**, decreased). The number of significantly changed proteins identified for each GO term is indicated in brackets. A single protein can be classified into multiple different GO terms. **(E)** Selected PPI clusters identified to be differentially regulated between fly brains with MB-specific *Atg5* knockdown and genetic controls. The size of individual nodes is defined relative to the corresponding −log_10_ p-value, and the color intensity of individual nodes indicates the size of the Log_2_ fold change between effect group and control. The thickness and intensity of individual strings between two nodes indicates the confidence score for a given interaction. The annotated GO terms were directly retrieved from Cytoscape.

Beyond ER-related signatures, remodeling of the presynaptic active zone emerged as a particularly prominent and specific feature of the proteomic response to MB-specific autophagy impairment. GO terms linked to presynaptic structure and function were significantly enriched across the differentially expressed protein set (Fig. 4A-D). Among the upregulated categories (Fig. 4A,C) were “presynaptic membrane” (GO:0042734), “presynaptic active zone organization” (GO:1990709), and “trans-synaptic signaling” (GO:0099537), whereas downregulated terms (Fig. 4B,D) included “voltage-gated calcium channel complex” (GO:0005891) and “modulation of chemical synaptic transmission” (GO:0050804). When focusing specifically on presynaptic proteins, we observed a robust and consistent upregulation of integral AZ components including BRP, RBP, RIM, Liprin-α, Syd-1, Unc13A, and Spinophilin (Spn) (Fig. 5A,C). In contrast, abundance of synaptic vesicle (SV)-associated proteins remained largely unaffected (Fig. 5B), and SV-related GO categories were coherently also not identified in the functional enrichment and PPI analysis (Fig. 4). These findings indicate that *Atg5* knockdown within the MB induces a selective and regulated restructuring of AZ protein composition, rather than a broad accumulation of synaptic proteins, supporting the interpretation that AZ remodeling is a hallmark of the PreScale-like resilience program.

**Fig. 5.**
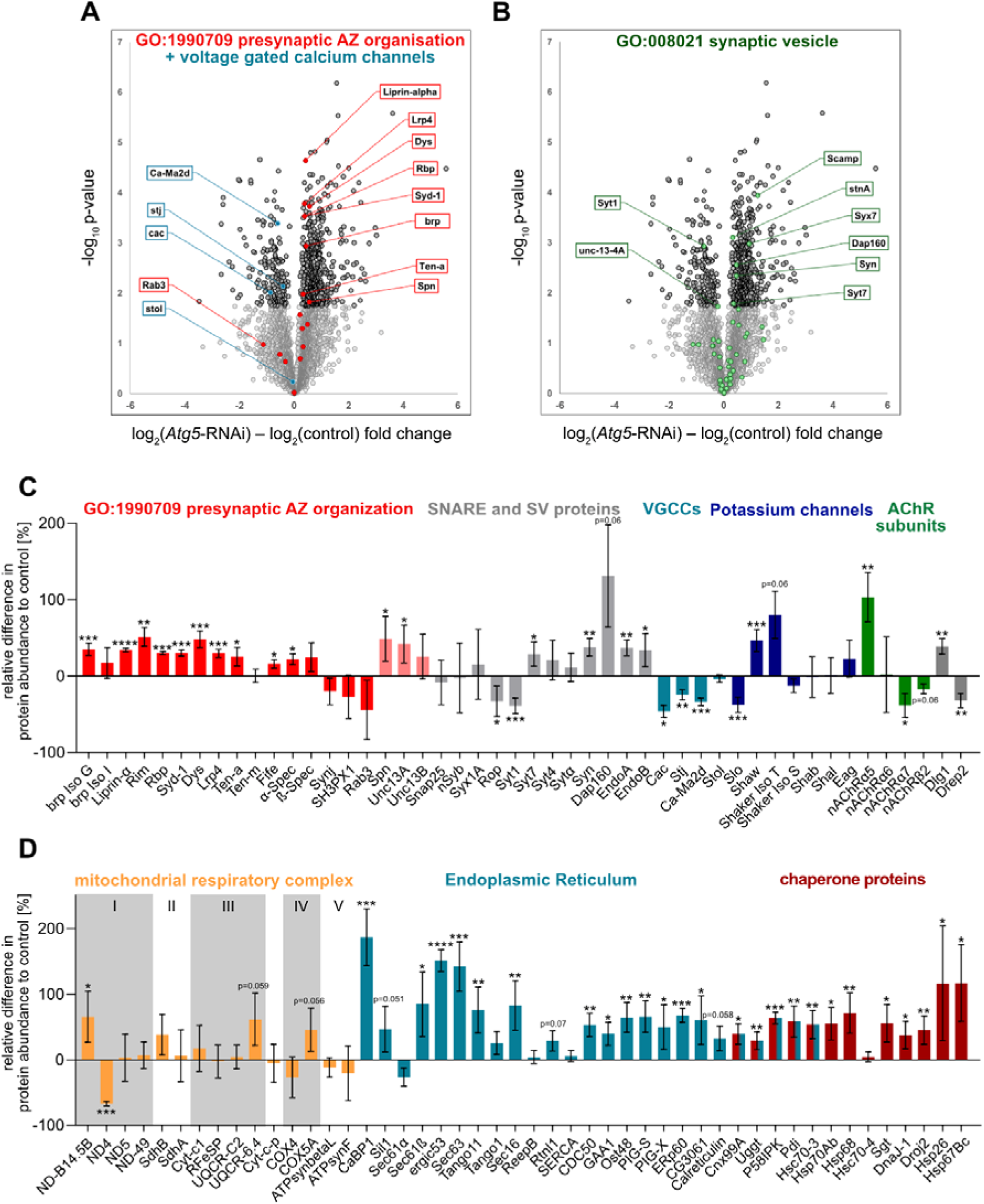
MB-specific *Atg5* knockdown provokes extensive changes in AZ and ER-related protein abundances. Selection of proteomic changes in brains of female *ok107*-Gal4>UAS-*Atg5*-RNAi flies compared to age- and sex-matched *ok107*-Gal4/+ controls. **(A)** Volcano plot of all proteins identified in the proteomic data set, highlighting candidates belonging to the GO term ‘presynaptic AZ organization’ (GO:1990709), as well as VGCCs components. The Log_2_ fold change on the x-axis is plotted against the −Log_10_ p-value of the two-tailed t-test comparison on the y-axis. **(B)** Volcano plot highlighting all identified proteins belonging to the GO term ‘synaptic vesicle’ (GO:008021). The Log_2_ fold change on the x-axis is plotted against the −Log_10_ p-value of the two-tailed t-test comparison on the y-axis. **(C)** Bar graphs of selected synaptic protein hits, including members of GO:1990709 ‘presynaptic AZ organization’ (red), additional AZ-associated proteins (light red), SNARE complex and synaptic vesicle (SV) components (light grey), VGCCs subunits (light blue), voltage-gated potassium channels (dark blue), acetylcholine receptor (AChR) subunits (green) and postsynaptic proteins (dark grey). **(D)** Bar graphs of selected protein hits, highlighting selected components of all 5 mitochondrial respiratory complexes (orange), ER-residing proteins (blue) and different classes of chaperone proteins (red). ER-specific chaperone proteins are indicated by 2 colors. For data presented in C and D, LFQ intensities measured for the effect group for each protein were normalized to the mean LFQ intensity of the control group to display differences in protein abundance upon MB-specific *Atg5* knockdown relative to the protein level of the control group. n = 3-4; two-tailed t-test; * = p<0.05; ** = p<0.01; *** = p<0.001; **** = p<0. 0001; Only significant differences are depicted. Data presented as mean ± SD.

However, not all presynaptic AZ components followed the same pattern. The voltage-gated calcium channel (VGCC) Cacophony (Cac) and its synapse-specific α2δ subunit straightjacket (stj) were significantly downregulated, while the dendritically localized α2δ subunit stolid (stol) remained unchanged (Fig-5C) ^55^. This selective reduction of axonal VGCC components points to a targeted reprogramming of presynaptic calcium signaling within axonal compartments, rather than a global reduction in channel expression. Such a shift is likely to decrease the probability of vesicle release in response to action potentials, tuning synaptic output in an activity-dependent manner. Along those lines, core SNARE proteins (Snap25, nSyb, Syx1A) remained unchanged (Fig. 5C), but the Ca² sensor Synaptotagmin-1 (Syt1) was moderately downregulated (Fig. 5B,C), reinforcing the idea of dampened release dynamics. On the other hand, several voltage-gated potassium channels, including Shaker (K_v_1) and Shaw (K_v_3), were robustly upregulated (Fig. 5C). Given the opposing effects of calcium and potassium conductance on presynaptic excitability — facilitating versus restraining neurotransmitter release respectively — these combined changes suggest an adaptive rebalancing of axonal excitability. Together, these findings point to a coordinated reprogramming of presynaptic terminals at the level of calcium influx and membrane excitability, likely contributing to the protective, low-gain transmission state characteristic of PreScale ^11^.

As already suggested by the GO functional enrichment and PPI analysis (Fig. 4), we also observed increased protein levels of numerous ER-related components in fly brains with MB-specific *Atg5* knockdown (Fig. 5D). Among the significantly increased hits were ER-components involved in protein folding (Calnexin99A (Cnx99a), Uggt, P58IPK, Hsc70-3, Pdi), GPI-anchoring (PIG-S), ER-organization (Sec16) and response to ER stress (Hsc70-3; CaBP1, Erp60, Uggt, Pdi) (Fig. 5D). Of note, we also observed increased levels of diverse classes of chaperone proteins including Hsp40/DNAj co-chaperones (DnaJ-1, Droj2), Hsp70 chaperones (Hsp68, Hsc70-3, Hsp70Ab) as well as small Hsps (Hsp26, Hsp67Bc) (Fig. 5D), indicating increased activity of protein-quality control systems upon impairment of autophagic degradation within the MB. Somewhat surprisingly, we could not detect major changes in mitochondria-specific proteins, suggesting that mitochondria might not be a major substrate of neuronal autophagy in this context. A lack of mitochondrial phenotypes had been reported also for murine neurons with conditional knockout of *Atg5* ^51^. Coherently, a directed search for proteins of all 5 complexes of the mitochondrial respiratory electron transport chain revealed no clear tendency towards increased or decreased abundances, with most components being unchanged compared to the control group (exemplary protein hits depicted in Fig. 5D). Furthermore, mitochondria-related GO terms were also not identified among the GO functional enrichment (Fig. 4A-D) and PPI network analyses (Fig. 4E).

### Extensive proteomic remodeling upon MB-autophagy impairment occurs largely independent of transcriptional changes

To determine whether these widespread proteomic changes originated at the transcriptional level or instead reflected post-transcriptional regulation, we performed RNA sequencing of fly heads from *ok107*-Gal4>UAS-*Atg5*-RNAi and *ok107*-Gal4/+ control flies. Strikingly, fewer than 6% of all significantly changed protein hits showed corresponding alterations at the transcript level (Extended Data Fig. 2). While some discrepancy could be due to technical factors, the data suggest that the majority of proteomic remodeling—including that of AZ proteins, VGCC subunits and ER components—is driven primarily by translational control or altered protein turnover, rather than changes in gene expression. Notably, however, two molecular chaperones — Hsp26 and Cnx99A — were transcriptionally upregulated (Extended Data Fig. 2) in parallel with their increased protein levels (Fig. 5D), consistent with a selective activation of protective protein quality control programs in response to MB-autophagy dysfunction.

### MB-specific *Atg5* knockdown results in brain-wide non-cell-autonomous accumulation of p62/ubiquitin aggregates often associated with ATG8a

ATG5 is an essential component of the autophagic machinery required for LC3/ATG8a lipidation, and thus is indispensable for phagophore expansion and autophagosome formation ^12,56–58^. In the *Drosophila* brain, pan-neuronal knockdown of *Atg5* leads to a complete loss of lipidated ATG8a-II and causes widespread accumulation of undegraded protein aggregates marked with the selective autophagy receptor Ref(2)P/P62 ^40^. In our PPI analysis of brain homogenates from MB-autophagy-impaired *ok107*-Gal4>UAS-*Atg5*-RNAi flies, we identified a small protein cluster annotated as ‘selective autophagy,’ containing mildly decreased ATG5 and strongly increased Ref(2)P/P62 levels (Fig. 4E). However, other autophagy-related GO terms and protein clusters were not identified by our analyses of significantly changed protein hits. A focused search for proteins belonging to the GO term ‘autophagy’ (GO:0016236) confirmed that multiple autophagy key components — including ATG4, ATG7, ATG12, and ATG8 — remained unchanged upon MB-specific *Atg5* knockdown compared to controls (Fig. 6A). Importantly however, Ref(2)P/P62 was identified as the most enriched protein in the *ok107*-Gal4>UAS-*Atg5*-RNAi group (Fig. 6A), indicating strong impairments of autophagic degradation and a drastic accumulation of undegraded protein aggregates.

**Fig. 6.**
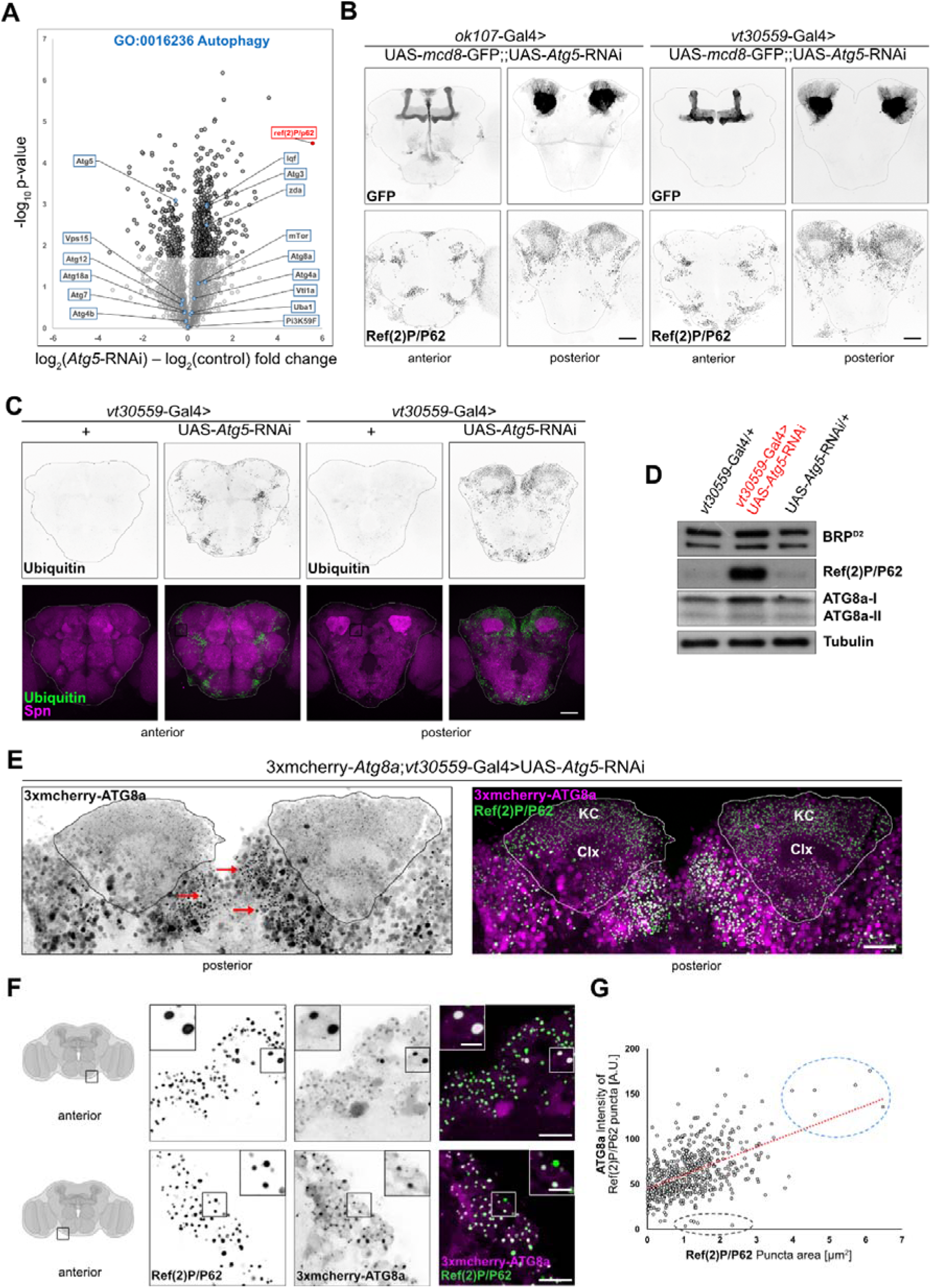
Widespread non-cell autonomous accumulation of autophagic substrates and ATG8a-positive structures upon MB-restricted *Atg5* knockdown. **(A)** Volcano plot highlighting all proteins belonging to the GO term ‘autophagy’ (GO:0016236) identified in the proteomic data set (Fig. 4 and Fig. 5). The Log_2_ fold change between *ok107*-Gal4>UAS-*Atg5*-RNAi effect and *ok107*-Gal4/+ control group is displayed on the x-axis, the −Log_10_ p-value of the two-tailed t-test comparison is plotted on the y-axis. **(B)** Representative confocal max projections of the anterior and posterior side of 5d old female fly brains simultaneously expressing a membrane-bound GFP (UAS-*mcd8*-GFP) and an RNAi against *Atg5* (UAS-*Atg5*-RNAi) under the control of either *ok107*-Gal4 or *vt30559*-Gal4. Whole-mount brain immunostaining was performed against Ref(2)P/P62 and BRP^Nc82^, and mcd8-GFP fluorescence was detected without additional antibody labelling. Scale bar = 50 µm. **(C)** Representative confocal max projections of the anterior and posterior side of 5d old female fly brains with MB-specific knockdown of *Atg5* (*vt30559*-Gal4>UAS-*Atg5*-RNAi) compared to a genetic control (*vt30559*-Gal4/+). Whole-mount brain immunostaining was performed against Ubiquitin and Spinophilin (Spn). Scale bar = 50 µm. **(D)** Representative Westernblot of whole brain homogenates of 5d old female flies with MB-specific knockdown of *Atg5* (*vt30559*-Gal4>UAS-*Atg5*-RNAi) and of the two genetic controls (*vt30559*-Gal4/+ and UAS-*Atg5*-RNAi/+). Membranes were probed with antibodies against BRP^D2^, Ref(2)P/P62 and ATG8a, differentiating between the unlipidated (ATG8a-I) and the autophagosome-associated lipidated (ATG8a-II) form. Tubulin was probed as loading control. The amount equal to two brains was loaded per lane. **(E)** Representative confocal max projections of the posterior side of the brain of a female *vt30559*-Gal4>UAS-*Atg5*-RNAi fly, immunostained with an antibody against Ref(2)P/P62 and simultaneously displaying endogenous 3xmCherry-ATG8a fluorescent signal. The MB KC regions (comprising the KC cell bodies (KC) and the Calyx neuropile (Clx)) are indicated by ROIs. Examples of non-cell autonomous 3xmCherry-ATG8a puncta outside the genetically targeted MB region are indicated by red arrows. Scale bar = 25 µm. **(F)** Representative confocal max projections of inferior brain regions close to the suboesophageal ganglion highlighting the non-cell autonomous accumulation of heterogenous Ref(2)P/P62 aggregates partially co-positive for 3xmCherry-ATG8a. Zoom-ins highlight large Ref(2)P/P62 puncta strongly co-positive for 3xmCherry-ATG8a (upper row), as well as a Ref(2)P/P62 punctum without 3xmCherry-ATG8a co-localization (lower row). Scale bar = 15 μm, zoom-in scale bar = 5 μm. **(G)** Plot of Ref(2)P/P62 puncta area in relation to the ATG8a intensity inside the Ref(2)P/P62 puncta. Quantification related to representative images in (F). The blue circle highlights the largest Ref(2)P/P62 puncta with high intensity mCherry-ATG8a co-labelling (exemplified in zoom-in in (F), upper row), grey circles indicate Ref(2)P/P62 aggregates without any mCherry-ATG8a co-label (exemplified in zoom-in in (F), lower row). n = 607 analyzed Ref(2)P/P62 puncta.

While a certain degree of Ref(2)P/P62 buildup was anticipated following MB-specific *Atg5* knockdown, the magnitude of this increase was surprising, given that only ∼3% of all neurons in the brain (approximately 4000 MB KCs) were genetically targeted. This was also reflected by the only mildly reduced total ATG5 protein levels in whole brain extracts (Fig. 4E,6A). To further investigate the spatial distribution of Ref(2)P/P62 accumulations across the brain, we used a UAS-*mCD8*-GFP transgene in combination with the UAS-*Atg5*-RNAi transgene driven by either *ok107*-Gal4 or *vt30559*-Gal4, to visualize Ref(2)P/P62 localization and abundance in relation to the genetically targeted and GFP-labelled MB neurons with reduced *Atg5* expression. Strikingly, numerous large Ref(2)P/P62-positive aggregates were located outside the GFP-labelled MB KCs in both anterior) and posterior brain areas (Fig. 6B), indicating a non-cell autonomous accumulation of undegraded protein aggregates and pointing towards either a propagation of autophagy dysfunction, or a spreading of autophagic substrates across the brain.

These findings were corroborated by an independent genetic approach, in which a UAS-*Cas9* transgene was expressed under *vt30559*-Gal4 control in flies ubiquitously carrying a guide-RNA to induce site-directed DNA double-strand breaks into the *Atg5* gene. This MB-restricted genome editing again resulted in widespread non-cell-autonomous Ref(2)P/P62 accumulation (Extended Data Fig. 3). As ubiquitination typically precedes Ref(2)P/P62-mediated cargo aggregation and degradation ^59–61^, we immunostained MB-autophagy impaired fly brains with an anti-ubiquitin antibody and observed numerous ubiquitin-positive puncta outside the MB in similar brain regions as the previously observed non-cell autonomous Ref(2)P/P62 aggregates (Fig. 6C), further confirming the ne accumulation of undegraded autophagic substrates across the brain.

Surprisingly, Western blot analysis of whole brain homogenates revealed that total ATG8a-II levels were not reduced in *vt30559*-Gal4>UAS-*Atg5*-RNAi flies compared to controls (Fig. 6D), suggesting that ATG8a lipidation, and thereby most likely also autophagosome formation, remained globally intact and reinforcing that the *Atg5* knockdown was indeed spatially confined to the MB KCs. Nonetheless, also a modest accumulation of unlipidated ATG8a-I was detected, consistent with cell autonomous effects of *Atg5* loss within MB KCs.

To better separate local (i.e., within the MB) from global, non-cell autonomous autophagy phenotypes, we combined *vt30559*-Gal4 and UAS-*Atg5*-RNAi fly lines with a ubiquitously expressed 3xmCherry-tagged endogenous *Atg8a* reporter ^62^. In the brain region surrounding the MB calyx where KC cell bodies reside, mCherry-ATG8a signal appeared diffuse and lacked punctate structures, despite the presence of numerous Ref(2)P/P62 aggregates (Fig. 6E), consistent with impaired autophagosome formation upon *Atg5* knockdown inside the MB. In contrast, increased, but still diffuse mCherry-ATG8a signal was detected within synaptic regions of the MB lobes (Extended Data Fig. 4), likely reflecting the accumulation of unlipidated ATG8a-I in presynaptic MB compartments, and eventually also explaining the increased level of ATG8a-I observed on Western blot level (Fig. 6D).

Outside the MB however, in neurons not targeted by *vt30559*-Gal4, we observed large and intense mCherry-ATG8a-positive puncta, which frequently co-localized with Ref(2)P/P62 aggregates and which could reach multiple µm in diameter (Fig. 6E,F). These ATG8a structures were particularly prominent in posterior regions between the calyces (Fig. 6E, red arrows), in anterior areas near the antennal lobes, and in inferior brain regions adjacent to the subesophageal ganglion (Fig. 6F). The co-localization of mCherry-ATG8a signal with Ref(2)P/P62 aggregates was highly heterogeneous: Some of the largest Ref(2)P/P62 puncta also displayed the highest intensity for mCherry-ATG8a (Fig. 6F,G, upper row zoom-in and blue circle), while others lacked clear mCherry-ATG8a co-labeling (Fig. 6F,G, lower row zoom-in and grey circle).

These findings support a model in which autophagy is disrupted cell autonomously upstream of autophagosome formation within MB KCs — consistent with the known requirement of ATG5 for ATG8 lipidation and autophagosome formation. In contrast, the widespread accumulation of Ref(2)P/P62 aggregates, co-labelled with ATG8a, in non-targeted neurons suggests that autophagosome formation remains functional outside the MB. This pattern indicates that while *Atg*5 knockdown directly blocks early autophagy steps in MB neurons, it also indirectly induces a secondary imbalance in autophagic degradation efficacy elsewhere in the brain. The heterogeneity of ATG8a recruitment to these aggregates further suggests regionally distinct compensatory responses to proteostatic stress, highlighting the complex nature of brain-wide autophagy propagation.

Taken together, these results demonstrate that MB-specific *Atg5* knockdown not only impairs autophagy locally but also leads to widespread, non-cell-autonomous accumulation of ubiquitinated and Ref(2)P/P62-labeled protein aggregates across the brain. The observed heterogeneity in mCherry-ATG8a recruitment suggests complex and spatially variable compensatory responses to local proteostatic stress.

## Discussion

Neural circuits must remain functionally stable while responding flexibly to changing demands, stressors, and aging-related decline ^63,64^. This balance is achieved through plasticity programs that integrate molecular, metabolic, and activity-dependent signals to reconfigure synaptic strength and connectivity ^65^. While many such programs are described at the level of individual synapses or pathways, much less is known about how localized stress responses — such as imbalances in proteostasis — can instruct circuit-wide adaptation to promote resilience. Autophagy, as a core mechanism for protein turnover and organelle quality control, is increasingly implicated in this form of adaptive plasticity.

This study reveals a molecularly resolved, brain-wide synaptic remodeling program triggered by local impairment of autophagy within the *Drosophila* mushroom body (MB). By combining targeted genetic manipulation, proteomic profiling, and circuit-level analysis, we identify a defined presynaptic adaptation characterized by active zone (AZ) scaffold expansion and altered excitability mediated by coordinated changes in calcium and potassium channels. These molecular changes emerge largely through post-transcriptional regulation and occur in the absence of widespread synaptic degeneration. Instead, they correlate with increased sleep, enhanced stress resilience, and extended survival — supporting the interpretation that they form part of a resilience-promoting plasticity program.

Previously, we described a non-cell autonomous increase in presynaptic abundance of the AZ scaffold protein BRP in response to MB-specific autophagy impairment ^40^, raising the possibility that local disruptions in proteostasis can instruct global synaptic remodeling. Here, we demonstrate that this effect is not limited to BRP and is embedded in a broader reorganization of presynaptic architecture. Our proteomic analysis revealed upregulation of several core AZ proteins — including RIM, RBP, Liprin-α, Syd-1, Unc13A, and Spinophilin (Spn), along with concurrent downregulation of the AZ VGCC components Cac and stj and upregulation of Shaker-type potassium channels (Fig. 4 + Fig. 5A,C). Together, these changes suggest a reconfiguration of release site number and vesicle release probability, consistent with a shift toward lower-gain, energetically efficient transmission ^11^.

This remodeling program was accompanied by a striking non-cell autonomous accumulation of protein aggregates labelled with the autophagy cargo receptor Ref(2)P/P62, both detected by proteomic approaches (Fig. 6A,D) and in spatially resolved immunohistochemical analyses using confocal microscopy (Fig. 6B + Extended Data Fig. 3). Notably, Ref(2)P/P62- and ubiquitin-positive aggregates accumulated throughout the brain — even in regions remote from the MB and without direct genetic manipulation — indicating a non-cell autonomous spread of autophagic stress or substrate buildup (Fig. 6). This observation points to circuit-level communication of proteostatic load, possibly via trans-synaptic or extracellular cargo transfer, and is consistent with findings in mammalian systems where protein aggregates and autophagic intermediates can spread between neurons or glia ^66–70^. Importantly, the widespread Ref(2)P/P62 aggregates in non-targeted brain areas were often associated with mCherry-ATG8a-positive structures, suggesting that autophagic degradation in these cells is dysbalanced downstream of autophagosome formation, at the level of cargo degradation efficiency (Fig. 6E-G). In contrast, within MB KCs where *Atg5* was knocked down, autophagy is blocked upstream, before autophagosome maturation, as expected based on ATG5’s essential role in ATG8 lipidation (Fig. 6E + Extended Data Fig. 4). This differential pattern of autophagic phenotypes supports the notion that local disruptions in autophagy can trigger a brain-wide, regionally heterogeneous breakdown in proteostatic handling, involving both direct and indirect mechanisms.

Importantly, our data show that this remodeling is not associated with overt functional decline. In contrast, MB-autophagy-impaired animals exhibit increased sleep (Fig. 2A-D), preserved oxidative stress resistance (Extended Data Fig. 1), and mild lifespan extension (Fig. 3C-H). These phenotypes align with the previously described PreScale plasticity — a brain-wide, transient presynaptic upscaling process marked by BRP elevation and linked to sleep need and early aging adaptation ^11,41^. Our experiments extend this framework by demonstrating that activation of PreScale-like remodeling can be accelerated via autophagy impairment, and conversely, can be delayed by preserving MB autophagy during early aging (Fig. 1). Specifically, enhanced BRP upregulation in MB *Atg5* knockdown animals was already detectable by 5 and 15 days of age (Fig. 1A-B), whereas *Atg5* overexpression suppressed brain-wide BRP increases in a similar time window around day 15 (Fig. 1E). By 30 days of age, differences between genotypes had levelled out (Fig. 1C,F), suggesting that MB-autophagy status controls the timing of PreScale activation but not its final extent, keeping levels of presynaptic AZ proteins in a physiologically relevant range (Fig. 1G).

This bidirectional control of PreScale kinetics positions the MB as a central node linking local metabolic and proteostatic stress to global circuit remodeling. The MB is well established as a hub for sleep regulation in Drosophila ^37,38^, integrating both sleep-promoting and wake-promoting pathways ^39,71^. Our findings suggest that it also acts as a proteostatic sensor, using its autophagy status to regulate presynaptic output and activity patterns in a manner that promotes restorative sleep and preserves resilience.

The causal link between synaptic remodeling and behavioral resilience is further supported by our genetic manipulation of PreScale levels in flies with MB-specific autophagy impairment. Decreasing *brp* gene dosage was sufficient to abolish the increase in survival observed in MB *Atg5* knockdown animals (Fig. 3I,J), underlining a functional role for BRP-driven AZ expansion in mediating these protective adaptive outcomes. These findings are in line with prior studies showing that BRP upscaling in PreScale is necessary for sleep homeostasis following deprivation ^41,43,72^ and sufficient to mediate sleep patterns changes in aging animals ^11^.

At the mechanistic level, our data suggest that MB autophagy status influences synaptic remodeling largely through post-transcriptional means. Despite pronounced changes in protein levels across AZ components and excitability regulators, corresponding transcript levels remained largely unchanged (Extended Data Fig. 2). This pattern points to regulated translation, protein turnover, or compartment-specific adjustments of proteostasis as key mechanisms for modulating synaptic protein composition. In this regard, the observed enrichment of ER-associated proteins and chaperones in our proteomic data is noteworthy (Fig. 4A,C + Fig. 5D). Although not the dominant feature of the remodeling program, ER remodeling and the unfolded protein response may support translational buffering and may contribute to the control of neurotransmission-relevant Ca^2+^ homeostasis at presynaptic compartments, as described in mammalian neurons ^51^.

Our findings also resonate with broader concepts in neurodegeneration and resilience. In mammalian systems, resilience to Alzheimer’s pathology has been linked to the preservation of specific synaptic protein modules ^73^, and autophagic flux is a key determinant of neuronal survival in both aging and disease ^24,25,74^. The MB, as a central integrator of synaptic, metabolic, and proteostatic information ^36^, may thus serve as a model for understanding how localized stress responses can be translated into brain-wide adaptive reprogramming.

In sum, we define a genetically tractable, mechanistically resolved program of presynaptic plasticity that links autophagy status in the MB to circuit-wide remodeling, sleep behavior, and stress resilience. This program shares key features with PreScale, including BRP-centered AZ expansion and behavioral restoration, but our data extend the model by identifying upstream triggers, molecular logic, and bidirectional regulatory control. More broadly, our work highlights the ability of local proteostatic stress to shape presynaptic architecture and function across the brain, establishing a framework for studying resilience as an emergent property of distributed synaptic plasticity.

The identification of a non-cell autonomous synaptic resilience program centered on presynaptic remodeling has important implications beyond basic neurobiology. Many neurodegenerative and neurodevelopmental disorders — including Alzheimer’s disease, Parkinson’s disease, and autism spectrum disorders — feature disrupted proteostasis, ER stress, and altered synaptic plasticity ^24,25,75,76^. The demonstration that local autophagy impairment can systemically reprogram synaptic function suggests new paths for understanding how focal cellular dysfunction spreads through neural networks. In particular, our work supports a model in which AZ remodeling acts as a compensatory adaptation to local autophagic stress, one that may initially preserve function but later become maladaptive. Elucidating this balance between resilience and pathological remodeling could open new therapeutic opportunities aimed at enhancing beneficial aspects of synaptic plasticity while curbing their long-term destabilizing effects.

**Extended Data Fig. 1.**
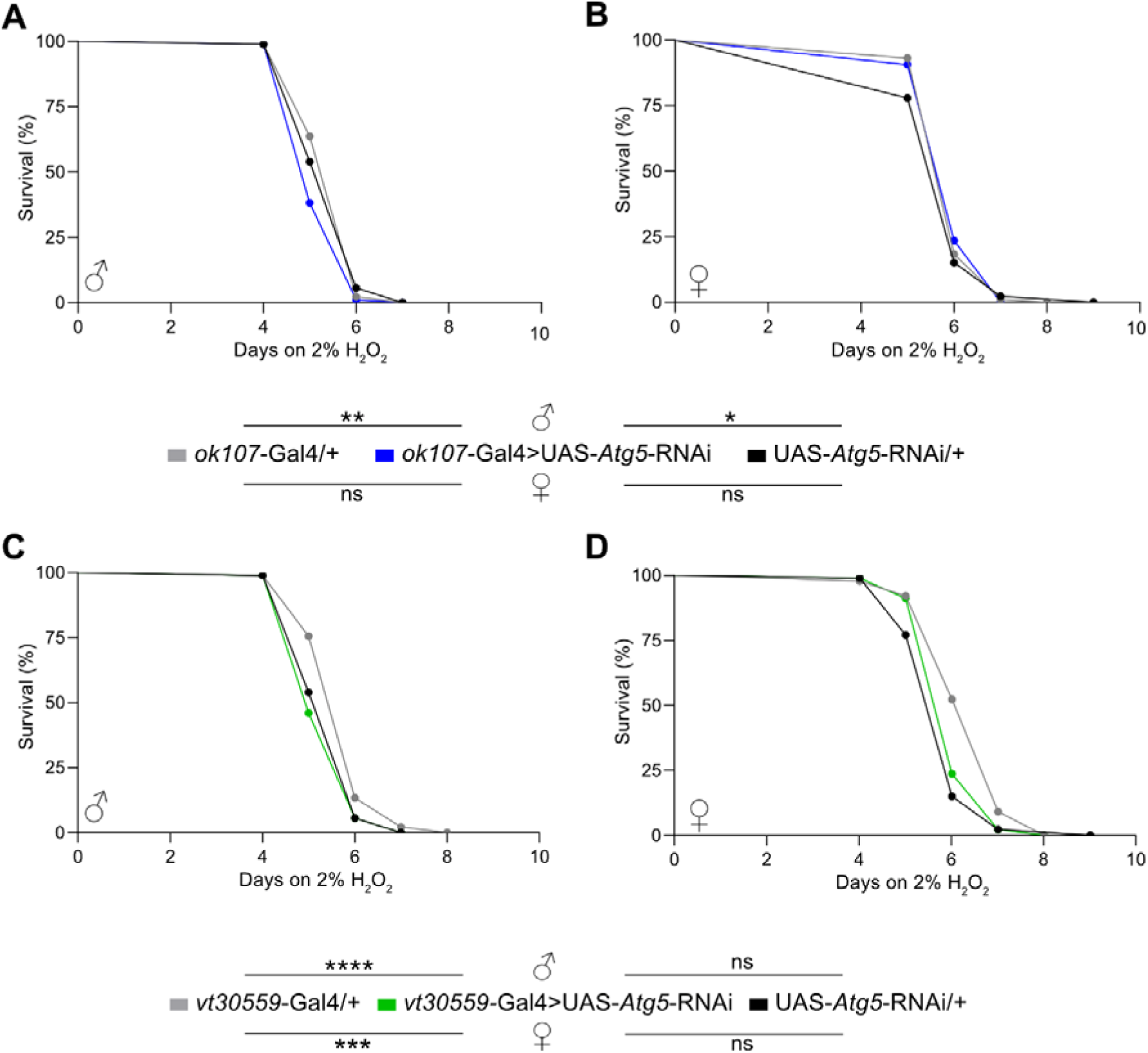
MB-specific *Atg5* knockdown does not attenuate H_2_O_2_ resistance. **(A and B)** Kaplan-Meier survival curves of male **(A)** and **(B)** female flies with MB-specific *Atg5* knockdown (*ok107*-Gal4>UAS-*Atg5*-RNAi, blue curve) compared to *ok107*-Gal4/+ and UAS-*Atg5*-RNAi/+ control flies continuously exposed to 2% H_2_O_2_ from day 2 onwards. n = 85-89 per genotype and sex. **(C and D)** Kaplan-Meier survival curves of male **(C)** and **(D)** female flies with MB-specific *Atg5* knockdown (*vt30559*-Gal4>UAS-*Atg5*-RNAi, green curve) compared to *vt30559*-Gal4/+ and UAS-*Atg5*-RNAi/+ control flies continuously exposed to 2% H_2_O_2_ from day 2 onwards. n = 87-90 per genotype and sex. The UAS-*Atg5*-RNAi/+ controls in (A) and (B) are the same as in (C) and (D). * = p<0.05; ** = p<0.01; *** = p<0.001; **** = p<0.0001; ns, not significant; pairwise Log-rank (Mantel-Cox) test comparison.

**Extended Data Fig. 2.**
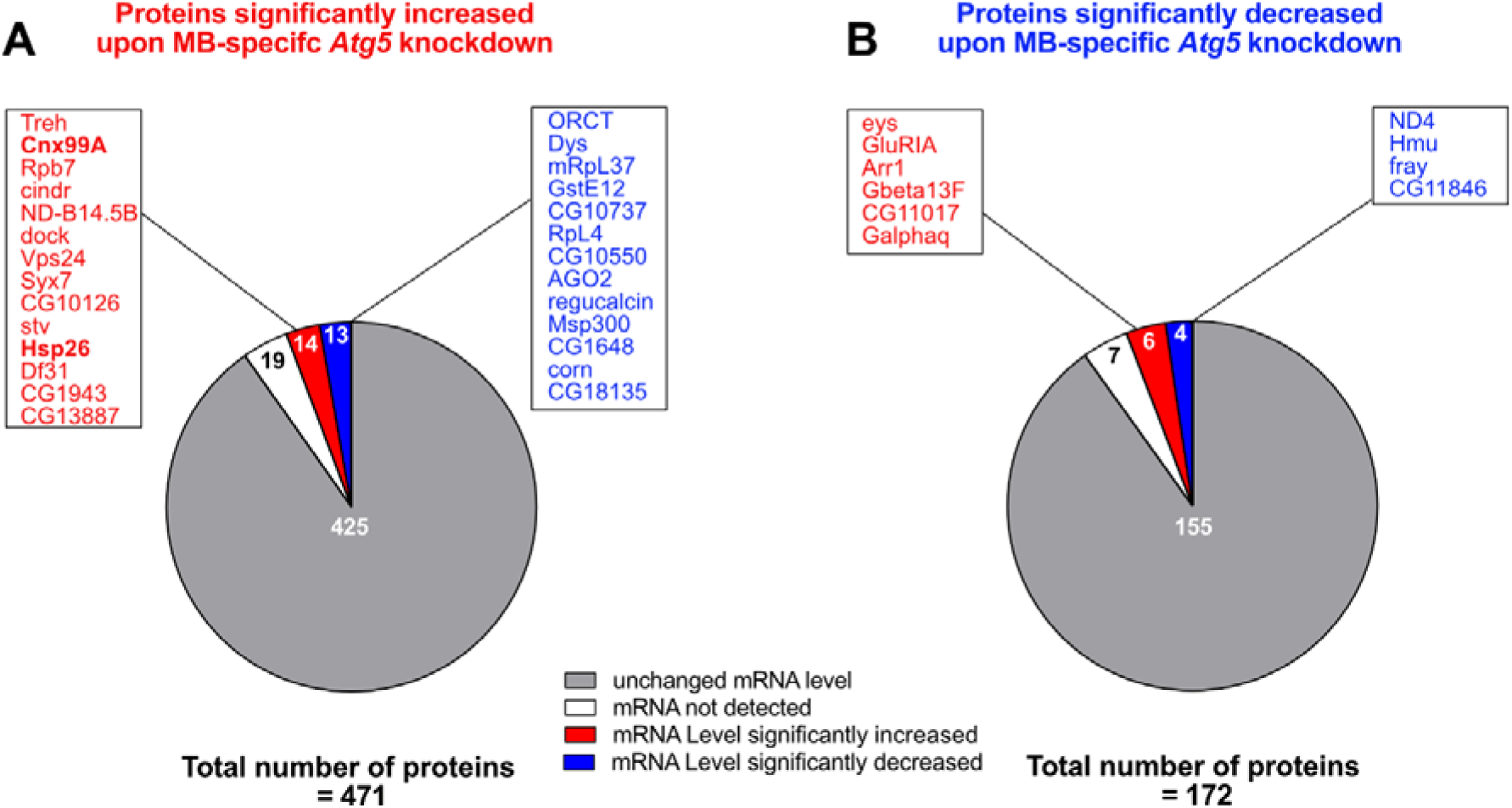
Proteomic changes in fly brains with MB-specific *Atg5* knockdown are largely mediated post-transcriptionally. Correlation of all significantly increased **(A)** or decreased **(B)** proteins in *ok107*-Gal4>UAS-*Atg5*-RNAi fly brains compared to *ok107*-Gal4/+ controls detected in the proteomic experiment with the data set acquired by RNA-Seq. Candidates with significantly changed expression at mRNA and protein level are depicted in the boxes.

**Extended Data Fig. 3.**
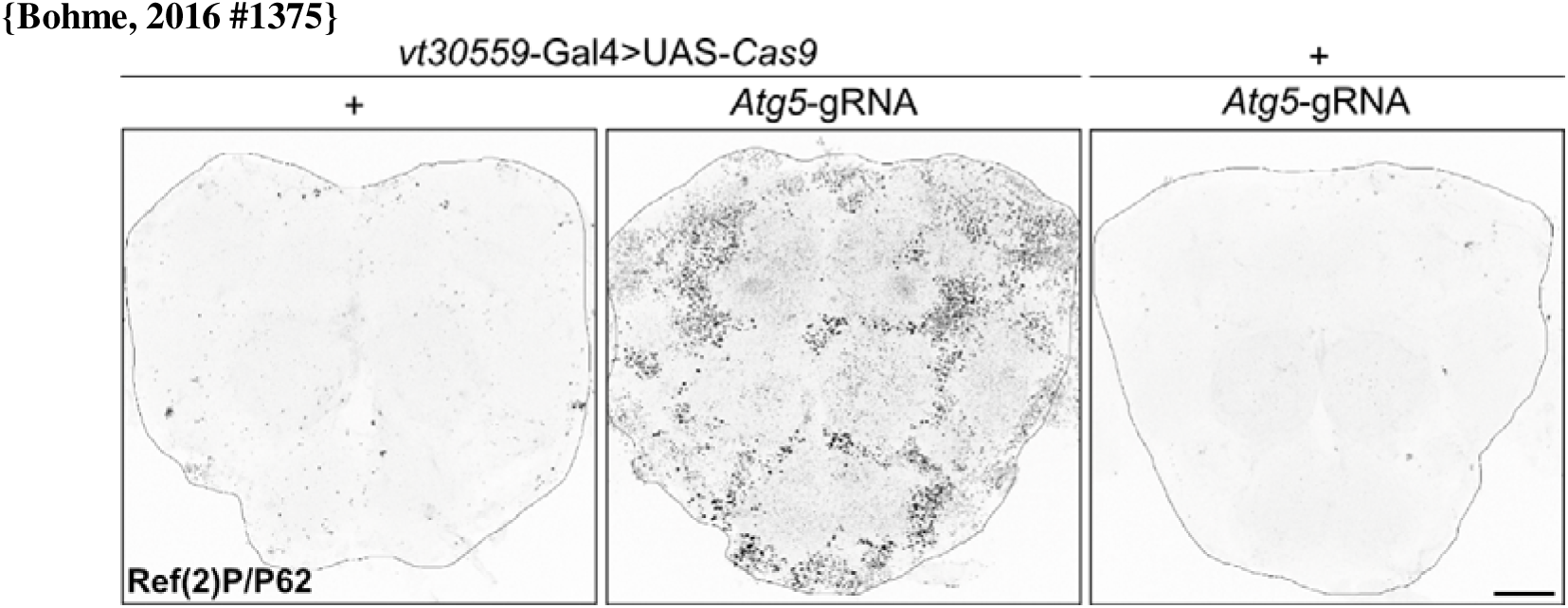
Somatic CRISPR/Cas9-mediated conditional knockout of *Atg5* specifically in MB KCs results in widespread non-cell autonomous accumulation of Ref(2)P/P62. Representative confocal max projections of whole-mount 5d old female fly brains immunostained against Ref(2)P/P62. The expression of *Cas9* in the MB under control of *vt30559*-Gal4 and the simultaneous presence of an ubiquitously expressed *Atg5*-gRNA induces site-directed DNA double stand breaks into the *Atg5* gene exclusively in the MB KCs. Scale bar = 50 µm.

**Extended Data Fig. 4.**
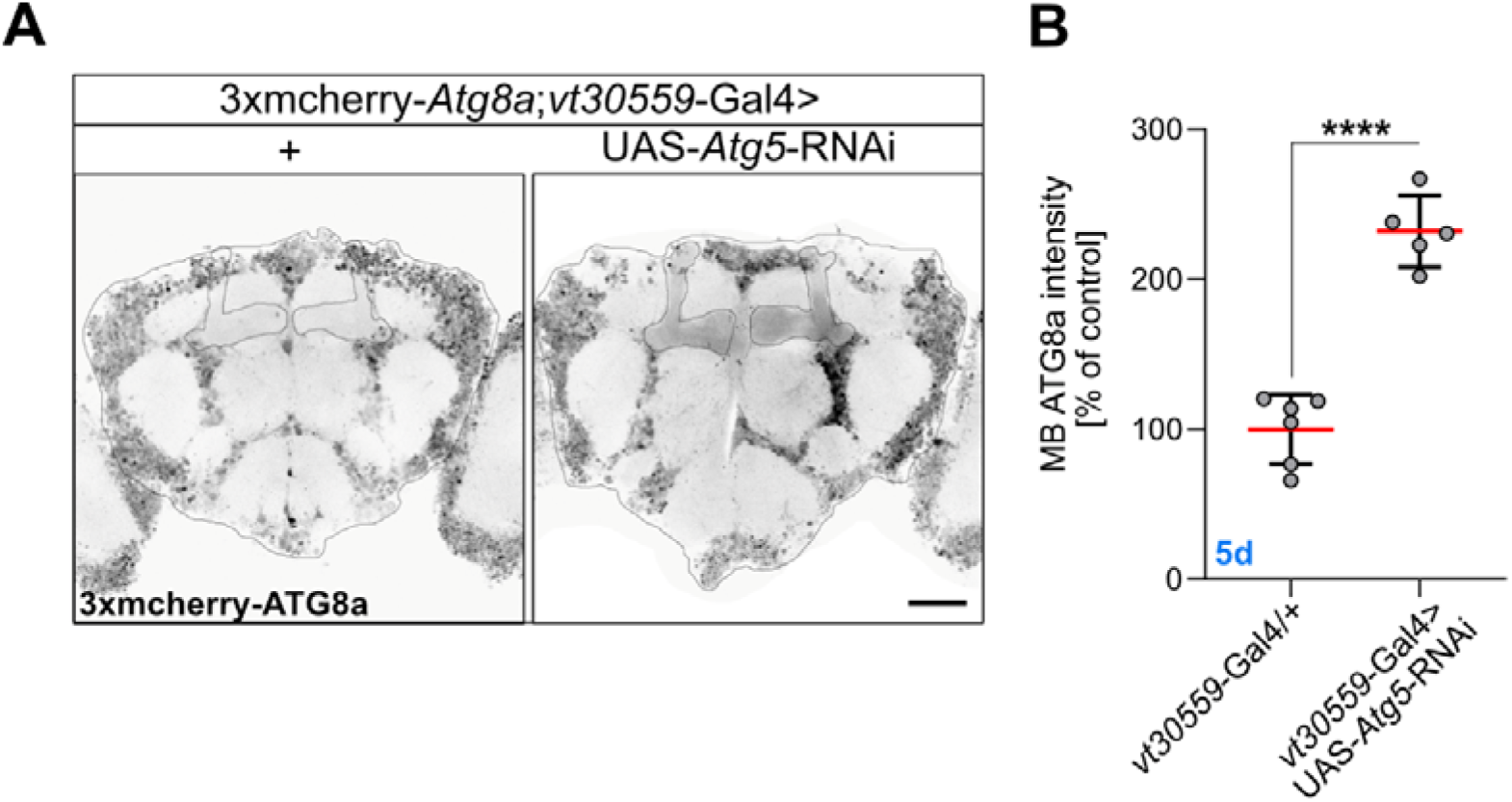
Synaptic accumulation of ATG8a in the MB lobes upon *vt30559*-Gal4 mediated knockdown of *Atg5*. **(A)** Representative confocal max projections of the fluorescent signal of the endogenous promoter-driven 3xmCherry-Atg8a reporter in vt30559-Gal4>UAS-Atg5-RNAi and vt30559-Gal4/+ control brains, imaged from the anterior side. Scale bar = 50 μm **(B)** Quantification of the 3xmCherry-ATG8a fluorescent signal inside the MB lobes. Representative ROIs surrounding the MB lobes are depicted in (A). n = 5-6. * = p<0.05; ** = p<0.01; *** = p<0.001; **** = p<0.0001; ns, not significant, two-tailed t-test. Data presented as mean ± SD.

## Material and Methods

### Drosophila stocks and maintenance

Fly stocks and experimental crosses were maintained under standard laboratory conditions on semi-defined food medium (Bloomington recipe) under a 12/12 h light/dark cycle with 65 % humidity at 25°C. Crosses and progeny for proteomics and RNA-Sequencing experiments (*ok107*-Gal4/+ and ok*107*-Gal4>UAS-*Atg5*-RNAi) were raised and maintained at 29°C ^40^. All flies were backcrossed to the same *w^1118^* (*iso31*, BDSC #5905) background for at least six generations, except for 3xmCherry-*Atg8a* flies. If not otherwise indicated, 5 days old flies were used for all experiments.

### Sleep measurements

Sleep and locomotor activity measurements of single adult flies was performed with *Drosophila* activity monitors (DAM2) from Trikinetics Inc. (Waltham, MA., USA) at 25°C and 65% humidity in a 12/12h light/dark cycle ^41,77^. At day 3 of age, individual flies were loaded under brief CO_2_ anesthesia into single Trikinetics glass tubes (5 mm inner diameter and 65 mm length) containing a food source of 5% sucrose and 2% agar on one side. The activity of individual flies was recorded as number of beam crosses within 1 min, and periods of inactivity without any beam cross of 5 mins or longer were considered as sleep ^78^. Activity measurements were performed over 5 consecutive days, and measurements from the first 2 days were discarded due to entrainment of the flies to the new environment. Data from the last 3 days was averaged and sleep parameters were analyzed using the Sleep and Circadian Analysis MATLAB Program (SCAMP) ^79^. Activity measurements of flies which died during the experiment were discarded from the analysis. For sleep measurements of aged flies, 13 days old flies were loaded into Trikinetics glass tubes so that they reached 15 days of age at the start of the relevant recording period.

### Longevity assay

Newly eclosed flies were collected every other day and maintained in groups of mixed sex for 2 days. At day 3, male and female flies were quickly separated under CO_2_ anesthesia and sorted into populations of approximately 25 individuals per vial. This procedure was repeated on different days with different cohorts of collected flies, until 6-10 experimental vials per sex and genotype were set up. The established longevity cohorts were transferred onto fresh food every second day, and the number of individual dead flies per vial was recorded at each timepoint of transfer.

### H_2_O_2_ survival assay

Freshly eclosed flies were collected every other day and maintained in groups of mixed sex for 2 days. Subsequently, 45 flies per genotype and sex were transferred onto big vials containing 3 filter papers soaked with 2 ml of solution consisting of 2% H_2_O_2_ and 5% sucrose diluted in water. Experimental cohorts were transferred onto new vials with freshly prepared H2O2 solutions every second day, and the number of individual death events per each vial was scored exactly at the same time of the day every 24 hours.

### Westernblot

Westernblot was performed as previously described ^80^, with minor modifications. In brief, 4 female adult brains per genotype were dissected in cold hemolymph-like saline solution (HL3 70 mM NaCl, 5 mM KCl, 10 mM NaHCO_3_, 20 mM MgCl_2_, 1.5 mM CaCl_2_, 5 mM HEPES, 5 mM Trehalose, 115 mM Sucrose, pH = 7.2) and stored in 50 µl HL3 in low-binding Eppendorf tubes on ice. Dissected brains were immediately centrifuged for 5 mins at 7000 rpm at 4°C, the HL3 supernatant was discarded, and the sample was frozen at −20°C. After completion of sample collection, 30 µl of Lysis buffer (1x PBS, 0.5% Triton-X, 2% SDS, 1x Sample Buffer, 1x Roche cOmplete™ Mini protease inhibitor cocktail (#11836153001)) was added onto the brain pellets, and samples were homogenized by mechanical rotation for 1 h at 4°C, followed by 5 mins full-speed centrifugation at 13.000 rpm at room temperature. All genotypes/groups of an experiment were dissected on the same day, and sample collection for Western blotting was always performed in the middle of the day (around ZT5, i.e., 2 pm). Lysed brain samples were boiled for 5 mins at 95°C and subsequently centrifuged for 1 min at 13.000 rpm. An amount equal to 2 brains for each experimental condition was loaded onto SDS-PAGE gels and immunoblotted according to standard protocols. Blots were manually developed with ECL solution and Kodak/GE films, and subsequently digitalized. The following primary antibodies were used: Rabbit anti-BRP^D2^ (rb5228 ^81^, 1:40.000), rabbit anti-Ref(2)P (Abcam Ab178440, 1:1000), rabbit anti-ATG8 (Abcam Ab109364, 1:500), mouse anti-Tubulin (Sigma T5168, 1:100.000). Secondary horseradish peroxidase (HRP)-conjugated goat anti-mouse and goat anti-rabbit antibodies were used for detecting the protein signals by chemiluminescence (Biozol, 1:5000).

### Sample preparation for label-free global proteomics of adult *Drosophila* brain homogenates

10 female brains per genotype were hand-dissected in cold HL3 and stored in 50 µl HL3 in low-binding Eppendorf tubes on ice, followed by centrifugation at 4°C and 7000 rpm for 5 mins, before being immediately frozen at −80°C until completion of sample collection. Dissection was always performed in the middle of the day (around ZT5, i.e., 2 pm), and the 4 biological replicates were dissected on different days from independently collected fly cohorts arised from the same parental cross. 20 µl of lysis buffer was added onto each frozen brain pellet, and samples were lysed for 1 hour at 4°C on a rotator, followed by 5 mins full-speed centrifugation at 13.000 rpm. Subsequently, 1 μl of 100 mM Dithiothreitol (DTT) dissolved in 50 mM Triethylammonium-Bicarbonat (TEAB) buffer was added to the samples to reach a final concentration of 5 mM of DTT in the solution, and samples were incubated shaking at 300 rpm for 30 mins at 55°C. Protein alkylation was performed by addition of 2.1 μl of 400 mM Chloracetematide (CAA) dissolved in 50 mM TEAB, to reach a final CAA concentration of 40 mM inside the sample. Samples were incubated shaking at 300 rpm for 30 mins at room temperature and in darkness, as CAA is instable under light exposure. After alkylation, proteomic samples were boiled for 5 mins at 95°C, followed by 1 min of centrifugation at 13.000 rpm. Exactly 20 μl of each sample was loaded onto 4-20% Mini-PROTEAN® TGX™ precast gels (Bio-Rad laboratories, Hercules, USA), and SDS-PAGE was performed for 2-3 mins at 200 V until all proteins migrated out of the loading pockets into the gel. After Coomassie staining of the gel, each lane of protein sample was cut with sterile scalpels into 4 equal pieces. Gel pieces were transferred into low-binding Eppendorf tubes containing 50 µl of distilled H_2_O, and stored at 4°C until further processing.

### Liquid chromatography mass spectrometry (MS) analysis

Tryptic digestion and MS analysis were performed at the Laboratory of Prof. Liu, FMP Berlin, Germany. In-Gel protein digestion was performed overnight at 37°C with trypsin at an enzyme to protein weight ratio of 1:20. Equal volumes corresponding to 1 μg of peptide were loaded on a Thermo Scientific Dionex UltiMate 3000 system connected to a PepMap C-18 trap-column (0.075 × 50 mm, 3 μm particle size, 100 Å pore size; Thermo Fisher Scientific), followed by an in-house-packed C18 column (column material: Poroshell 120 ECC18, 2.7 μm; Agilent Technologies). After loading, peptides were separated at 250 nL/min flow rate over a 120 min gradient of increasing acetonitrile concentration and sprayed into an Orbitrap Fusion Lumos (instrument control software 3.1). The MS1 scans were performed in the Orbitrap. Only precursors with a charge state of 2–4 were subjected to fragmentation and then dynamically excluded for 60 s. The MS2 scans were acquired in the ion trap with the following settings: scan rate rapid, 35 ms max. injection time, first mass 110 m/z, isolation window 1.6 m/z, 30% NCE, AGC target 10,000. A 1 s cycle time was set between master scans.

### Proteomic data analysis

#### Database search, data processing and statistical analysis

Raw data was searched using MaxQuant version 1.6.2.6 in default settings. The number of missed tryptic cleavages allowed was set to 2, ‘label-free quantification’ was enabled, and the‘match between runs’ option was disabled. The search was performed using the UniprotKB database of *Drosophila melanogaster* downloaded in May 2020 and containing 42,678 entries. Peptide-spectrum match (PSM) and protein false-discovery rate (FDR) were set to 1%. Further data analysis was performed with Perseus (v1.6.2.3). Label-free quantification (LFQ) values were log_2_ transformed to achieve a normal data distribution. Proteins identified in at least three (out of four) biological replicates were considered for further statistical quantification. Proteins that were detected and quantified in only one replicate were excluded. Missing data values were imputed by values from a normal distribution (width 0.3 standard deviations; down shift 1.8). For statistical protein enrichment analysis, the two experimental groups (i.e., o*k107*-Gal4>UAS-*Atg5*-RNAi and *ok107*-Gal4/+) were compared by unpaired two-tailed t-test with a significance level α = 0.05. Fold changes were calculated as the difference of mean log_2_ transformed intensity values between *ok107*-Gal4>UAS-*Atg5*-RNAi effect and *ok107*-Gal4/+ control group.

### Gene Ontology (GO) functional enrichment analysis

GO functional enrichment analysis was performed with Metascape ^54^. All identified significantly up- or downregulated proteins were uploaded to the Metascape website (https://metascape.org/), and functional enrichment analysis was performed separately for GO ‘Biological processes’ and GO ‘cellular components’ with a p-value cutoff of p<0.01, an enrichment factor >1.5, and a number of hits ≥2.

### Protein-Protein Interaction (PPI) network analysis

PPI network analysis was performed with Cytoscape (https://cytoscape.org/, Version 3.8.2). Log_2_ fold change (FC) and −log_10_ p-values of all significantly changed proteins were loaded into the software, and a full string network with a confidence score of 0.4 and 0 additional interactors was created. The size of individual nodes was defined relative to the corresponding −log_10_ p-value, and the color intensity of individual nodes indicates the size of the Log_2_ FC. The size and intensity of the strings between two nodes indicate the confidence score for a given interaction.

### RNA-Sequencing

Sample collection for RNA-Sequencing was performed as follows: Female flies of the desired age and genotypes were snap-frozen in liquid nitrogen and immediately stored at −80°C. Subsequently, frozen flies were decapitated with metal sieves without interrupting the cooling chain, and fly heads were immediately transferred into low-binding Eppendorf tubes to be stored at −80°C. In total, 4 biological replicates per genotype with approximately 50 heads/replicate were collected, and QuantSeq 3’mRNA sequencing was subsequently performed by Lexogen® (Vienna, Austria). Differential expression analysis was provided by the company and was performed with the DESeq2 Bioconductor implemented in R. Significantly changed gene expression between *ok107*-Gal4>UAS-*Atg5-*RNAi effect and *ok107*-Gal4/+ control group was defined by an adjusted p-value<0.10.

### Immunohistochemistry of adult *Drosophila* brains

Immunohistochemical stainings of adult *Drosophila* brains was performed as previously described ^40^ with minor modifications. In brief, flies with the desired age, sex and genotype were anesthetized on ice, brains were hand-dissected in 50 μl HL3 and stored in glass staining chambers containing 500 μl of HL3 on ice. Brains were immediately fixed in 4% PFA for 40 min at room temperature on a horizontal shaker. After fixation, brains were quickly rinsed 2 times, followed by 3 times of 10 min washing in 0.7% PBS-T. Subsequently, brains were incubated for 2 hours at room temperature shaking in 0.7% PBS-T containing 10% Roti® ImmunoBlock (Carl Roth GmbH). Brains were incubated gently shaking for 48 hours at 4°C in 200 µl of solution containing primary antibodies diluted in 0.7% PBS-T with 5% Roti® ImmunoBlock. In the following, brains were washed in 30 mins intervals at least 8 times with 0.7% PBS-T, and then incubated with corresponding fluorophore-conjugated secondary antibodies diluted in 0.7% PBS-T containing 5% Roti® ImmunoBlock overnight at 4°C on a shaker. Brains were washed in 30 mins intervals at least 6 times with 0.7% PBS-T, and subsequently mounted on glass slides using Vectashield® Mounting Medium (Vector Laboratories Inc.). Samples were covered with high-precision coverslips No. 1.5H (Carl Roth GmbH) and sealed with nail polish. Mounted brains were stored at −20°C in darkness until the day of microscopy image acquisition to preserve the fluorescence. The overview and exact dilutions of antibodies used for immunohistochemistry can be found in table 1.

### Confocal image acquisition

Confocal images of immunofluorescently stained adult *Drosophila* brains were acquired with a Leica TCS SP8 inverted confocal microscope (Leica Microsystems GmbH) at the BioSupraMol Optical Microscopy Facility of the Freie Universität Berlin, Germany. Optical slices of the whole brain were acquired at a format of 1024 x 1024 pixels and 0.75x zoom with a 40x, 1.3 NA (numerical aperture) oil immersion objective. Image stacks were acquired in line scanning mode at a speed of 400 Hz and at a Z-step size of 1 μm. For higher magnifications, selected brain areas were scanned with a 63x, 1.4 NA oil immersion objective and variable zoom and Z-step size settings, while all other parameters remained the same as for whole brain image acquisition. All settings were kept exactly the same for the image acquisition of all groups/conditions of a given experiment.

### Confocal image processing and analysis

Image, processing, analysis and the selection of representative stack projections was performed with Fiji/Image J software (Version 1.53q; ^82^).

### Whole brain mean fluorescence intensity measurement

Measurements of whole brain mean fluorescence intensities was performed on average projections of confocal image stacks, comprising all optical slices between the beginning of the antennal lobes at the anterior side and the end of the dorsal fan-shaped body brain structure towards the posterior side. A region of interest (ROI) was manually drawn around the whole brain and the mean grey value inside the ROI was measured. Individual intensities of all groups of a given experiment were normalized to the mean intensity of the control group in order to obtain the relative difference between control and experimental groups. For whole brain BRP intensity measurements, each experiment was independently performed twice, and the normalized intensity values obtained for each experiment were pooled together for analysis. For analysis of 3xmCherry-ATG8a intensities inside the MB lobes, a ROI was manually drawn around the MB lobes for single optical slices at an interval of 4 slices. Subsequently, missing ROIs were interpolated and manually corrected if needed, and 3xmCherry-ATG8a intensity was measured for each ROI per individual optical slice to obtain an average intensity value over the whole MB neuropile. Measured values were subsequently normalized to the average value of the control group.

### Ref(2)P/P62 and 3xmCherry-ATG8a puncta analysis

Quantification of Ref(2)P/P62 puncta number, size and intensity was performed on single optical slices. Images were thresholded and ROIs for individual Ref(2)P/P62 puncta were obtained with the ‘Find Maxima’ and ‘Analyze particles’ functions in FIJI/ Image J. The same ROIs obtained from Ref(2)P/P62 puncta were applied to the 3xmCherry-ATG8a channel, in order to obtain the corresponding 3xmCherry-ATG8a intensity for individual Ref(2)P/P62 puncta.

### Statistical analysis and visual data presentation

Data sets were analyzed and plotted into representative graphs using GraphPad Prism 8 (GraphPad Software Inc.). Statistical comparisons between two experimental groups was performed by unpaired two-tailed t-test, and statistical testing between more than two experimental groups was achieved by one-way analysis of variance (ANOVA) followed by Tukey’s *post-hoc* test for multiple comparisons. Kaplan-Meier-survival curves were statistically compared pairwise by Log-rank (Mantel-Cox) Test. The linear regression model for Fig. 6G was calculated in Excel (Microsoft). Statistical significance is indicated as follows: * = p<0.05; ** = p<0.01; *** = p<0.001; **** = p<0.0001; ns, not significant = p>0.05. All figures were created with Affinity Designer 2.5.6 (Serif).

## Resources

**Table 1:**
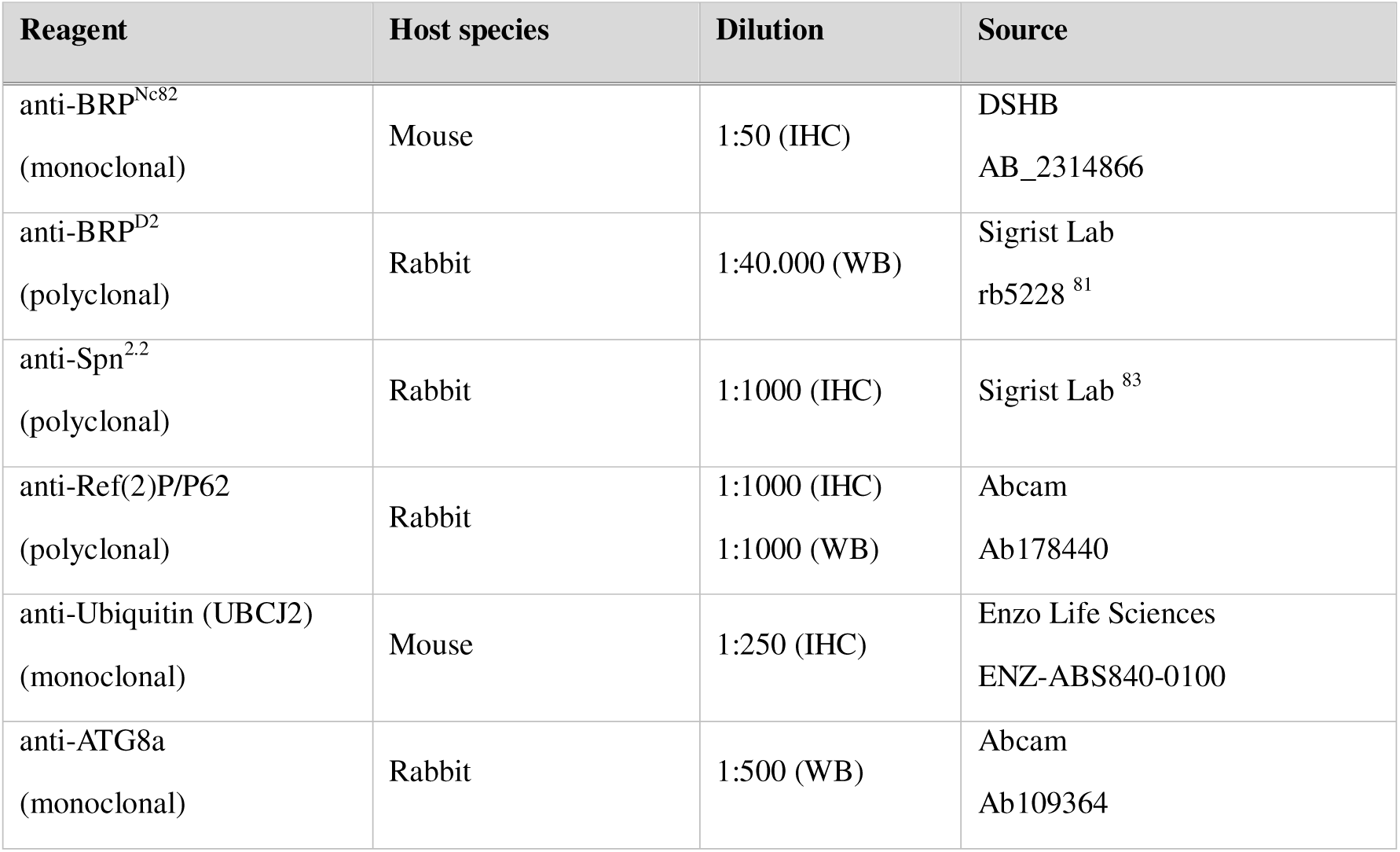
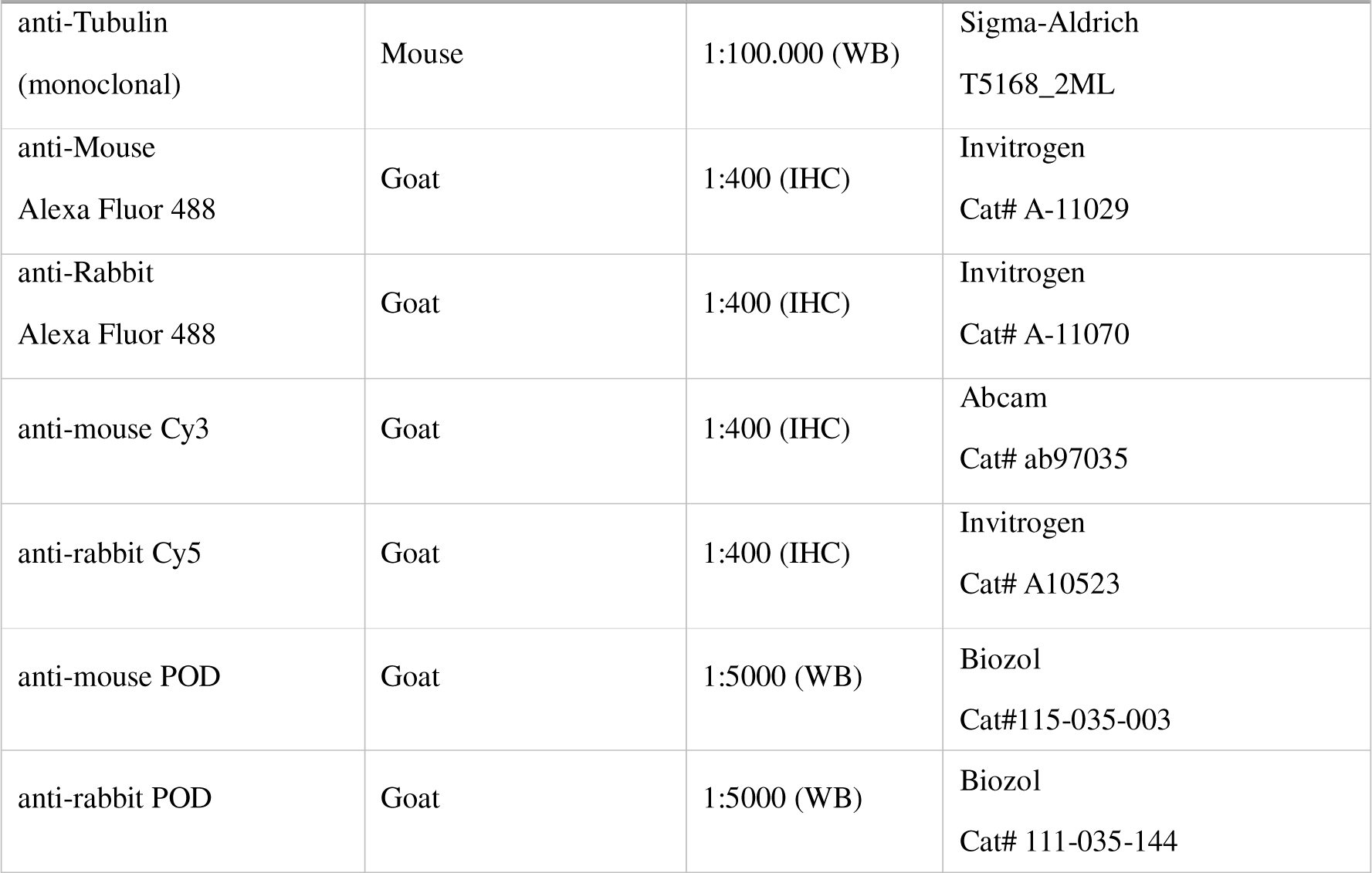
Antibodies.

**Table 2:** Biological samples.

| Designation | Source | Identifier |
| --- | --- | --- |
| $w^{1118}$ (Wild type) | Bloomington *Drosophila* Stock Center | #5905 |
| $Atg7^{d77}$ | <sup>47</sup> | N/A |
| $sss^{P1}$ | Bloomington *Drosophila* Stock Center | #16588 |
| 1x BRP | Bloomington *Drosophila* Stock Center | #85966 |
| 3xmCherry-*Atg8a* | <sup>62</sup> | N/A |
| *elav*-Gal4 | Bloomington *Drosophila* Stock Center | #458 |
| *vt30559*-Gal4 | Vienna *Drosophila* Resource Center | #206077 |
| *ok107*-Gal4 | <sup>52</sup> | N/A |
| UAS-*Atg5*-RNAi | Bloomington *Drosophila* Stock Center | #34899 |
| UAS-GFP-*Atg5* | Bloomington *Drosophila* Stock Center | #59848 |
| UAS-*Atg7* | <sup>84</sup> | N/A |
| UAS-*mcd8*-GFP | Sigrist Lab | Stock #15 |
| UAS-*Cas9* | Bloomington *Drosophila* Stock Center | isolated from<br>#67073 |
| *Atg5*-gRNA | Bloomington *Drosophila* Stock Center | #80819 |

## Acknowledgements

We thank Bloomington *Drosophila* Stock Center (BDSC) and Vienna *Drosophila* Resource Center (VDRC) for fly lines. We thank Gábor Juhász for the 3xmcherry-*Atg8a* line and for fruitful discussions. This work was supported by the European Commission (ERC Advanced Grant “SynProtect”: #101097053 to S.J.S.), and by grants from the Deutsche Forschungsgemeinschaft to S.J.S.: FOR5228 (#447288260).

## Author contributions

Conceptualization: D.T., S.H., S.J.S.; Investigation: D.T., S.H., J.L., F.L., M.M.; Funding acquisition and supervision: S.J.S.; Writing: S.J.S.;

